# Large-scale Phenotypic Drug Screen Identifies Neuroprotectants in Zebrafish and Mouse Models of Retinitis Pigmentosa

**DOI:** 10.1101/2020.03.26.010009

**Authors:** Liyun Zhang, Conan Chen, Jie Fu, Brendan Lilley, Cynthia Berlinicke, Baranda Hansen, Ding Ding, Guohua Wang, Tao Wang, Daniel Shou, Ying Ye, Meera T. Saxena, Kelsi R. Hall, Abigail V. Sharrock, Carlene Brandon, Joong Sup Shim, Justin Hanes, Hongkai Ji, Jun O. Liu, Jiang Qian, David F. Ackerley, Baerbel Rohrer, Donald J. Zack, Jeff S. Mumm

## Abstract

Retinitis pigmentosa (RP) and associated inherited retinal diseases (IRDs) are caused by rod photoreceptor degeneration, necessitating therapeutics promoting rod photoreceptor survival. To address this, we tested compounds for neuroprotective effects in zebrafish and mouse RP models, reasoning drugs effective across species may translate better clinically. We first performed a large-scale phenotypic drug screen using a larval zebrafish model of inducible RP. 2,934 compounds, mostly human-approved drugs, were tested across six concentrations. Statistically, 113 compounds achieved “hit” status. Secondary tests of 42 high-priority hits confirmed eleven lead compounds. Nine leads were then evaluated in mouse RP models, with six exhibiting neuroprotective effects. An analysis of potential mechanisms of action suggested complementary activities. Paired lead compound assays in zebrafish showed additive neuroprotective effects for the majority. These results highlight the value of cross-species phenotypic drug discovery and suggest combinatorial drug therapies may provide enhanced therapeutic benefits for patients with RP and IRDs.

## INTRODUCTION

Inherited retinal diseases (IRDs) encompass a group of genetically-linked retinopathies characterized by progressive photoreceptor death (Duncan et al., 2018). IRDs lead to irreversible vision loss, for which treatment strategies are limited. Retinitis pigmentosa (RP), the most common IRD, is characterized by early onset night blindness, gradual loss of visual field, and eventual loss of central vision (Ferrari et al., 2011; Hamel, 2006). Approximately 1.5 to 2.5 million RP patients are affected worldwide (Dias et al., 2017; Hartong et al., 2006; Verbakel et al., 2018). The initial pathological feature is selective rod photoreceptor cell death, which is generally followed by loss of cone photoreceptors (Léveillard et al., 2014). Mutations in more than 70 genes have been linked to RP (Dias et al., 2017; https://sph.uth.edu/retnet/). How these mutations affect gene function or initiate aberrant photoreceptor cell loss is largely unknown.

As RP/IRD progression is relatively protracted, pharmacological interventions aimed at slowing photoreceptor death are sought (Duncan et al., 2018; Wubben et al., 2019). However, currently there are no effective therapies for promoting photoreceptor survival. As a means of discovering new pharmacological treatments, target-directed high-throughput screening (HTS) approaches have been highly successful in identifying compounds that bind to and/or modulate disease-implicated molecules. However, many promising leads fail during late-stage animal model testing or clinical trials (Munos, 2009; Sams-Dodd, 2013; Scannell et al., 2012). This trend has renewed interest in phenotypic drug discovery (PDD), a complementary approach where drug effects are evaluated in cells or living disease models (Bickle, 2010; Lee et al., 2012; Swinney, 2013). A number of first-in-class drugs were recently discovered using PDD (Eder et al., 2014; Swinney and Anthony, 2011; Swinney, 2014). To expand opportunity on this front, we developed a PDD platform enabling quantitative HTS (qHTS; Inglese et al., 2006) in zebrafish (Walker et al., 2012; G. Wang et al., 2015; White et al., 2016).

Zebrafish offer several distinct advantages as a retinal disease modeling system (Angueyra and Kindt, 2018; Richardson et al., 2017). First, the structure of the zebrafish retina is similar to the other vertebrates (Angueyra and Kindt, 2018; Richardson et al., 2017; Schmitt and Dowling, 1999). In particular, the zebrafish retina is “cone rich” like the human retina. Second, about 70% of human genes have at least one ortholog in zebrafish (Howe et al., 2013). Moreover, all RP-associated genes listed in RetNet (https://sph.uth.edu/retnet/) have conserved zebrafish orthologs. Third, the zebrafish retinal system develops quickly being fairly mature by day five of development (Brockerhoff et al., 1995; Moyano et al., 2013; Schmitt and Dowling, 1999). Fourth, zebrafish are amendable to large-scale chemical screening due to their high fecundity rate, small size, and ease of visualizing and quantifying a variety of phenotypes (Mathias et al., 2012; Zon and Peterson, 2005). To streamline such screens, we developed a high-throughput plate reader-based method for quantifying reporter gene expression *in vivo* (‘ARQiv’; Walker et al., 2012). Recently, we adapted the ARQiv system to human stem cell-derived retinal organoids (Vergara et al., 2017) to enable cross-species PDD. To realize full throughput potential, ARQiv was combined with robotics-based automation to create “ARQiv-HTS” (G. Wang et al., 2015; White et al., 2016).

Here, to identify neuroprotective compounds promoting rod photoreceptor survival, ARQiv-HTS was used to perform a large-scale chemical screen in an inducible zebrafish model of RP (Walker et al., 2012; White et al., 2017). Close to 3,000 largely human-approved drugs were tested across six concentrations (i.e., using qHTS principles, Inglese et al., 2006) in more than 350,000 zebrafish larvae. Statistically, 113 hits were identified as hits and 42 of the top performing compounds advanced through validation and orthogonal assays. Eleven compounds passed all secondary tests and moved forward as lead compounds. Subsequently, a subset of leads was tested in primary mouse retinal cell cultures and retinal explants from retinal degeneration 1 mutant mice *(Pde6b^rd1^*, hereafter *rd1*), an RP model. Six leads showed neuroprotective effects in at least one mouse assay, and two showed activity in all retinal culture assays. One of these, dihydroartemisinin (DHA), was formulated for long-term release and evaluated *in vivo* using a second RP model, retinal degeneration 10 (*Pde6b ^rd10^*, hereafter *rd10*), but failed to show neuroprotection. Lastly, an analysis of potential mechanisms of action (MOA) of the eleven lead compounds suggested possible complementation. We therefore tested pairs of lead compounds for additive neuroprotective effects in zebrafish. Intriguingly, additive effects were evident for the majority of pairs tested. This result suggests combinatorial therapeutics developed with these reagents may provide enhanced neuroprotective effects. We posit that drugs able to function as neuroprotectants across diverse model species, and/or through complementary mechanisms, provide promising new therapeutic opportunities for RP/IRD patients.

## RESULTS

### Establishing a large-scale neuroprotectant screen using an inducible zebrafish RP model

The ARQiv-HTS platform was used to screen compounds for neuroprotective effects in a transgenic zebrafish model of RP, *Tg(rho:YFP-Eco.NfsB)gmc500*, hereafter, *rho:YFP-NTR* (Walker et al., 2012; White et al., 2017). In this line, a 3.7 kb *rhodopsin* (*rho)* promoter fragment (Hamaoka et al., 2002) drives transgene expression exclusively in rod photoreceptor cells. The transgene is a fusion protein linking a yellow fluorescent protein (YFP) reporter to a nitroreductase prodrug converting enzyme (NTR, encoded by the *E. Coli nfsB* gene). NTR expression enables pro-drug inducible targeted cell ablation (Curado et al., 2007). Exposing *rho:YFP-NTR* fish to the prodrug metronidazole (Mtz) leads to the selective death of rod photoreceptors and concomitant loss of YFP (Supplementary Fig. 1), modeling the onset of RP (Arango-Gonzalez et al., 2014; Chaitanya et al., 2010; Ripps, 2002; Sancho-Pelluz et al., 2008). To identify neuroprotective compounds that promote rod photoreceptor survival (i.e., sustain YFP expression after Mtz exposure) our plate reader-based ARQiv assay was used.

We first determined optimal conditions for inducing rod cell loss while maintaining larval health in a 96-well plate format. Major aspects of retinal cytogenesis are largely complete by 5 days post-fertilization (dpf) in zebrafish (Schmitt and Dowling, 1999; Stenkamp, 2011). Reporter expression in *rho:YFP-NTR* larvae has also stabilized by this time point (Unal Eroglu et al., 2018), consistent with *rho* expression (Raymond et al., 1995). We therefore chose 5 dpf to initiate Mtz-induced rod cell ablation. We previously determined that rod cell loss reached a nadir at 7 dpf following a 24 hr pulse of 10 mM Mtz at 5 dpf (Walker et al., 2012). We reasoned that a 48 hr Mtz exposure initiating at 5 dpf would maximize the signal window to test for neuroprotective effects. Concluding the experiment by 7 dpf also avoids challenges associated with feeding, as zebrafish can subsist on their yolk sac up to that time point (Hernandez et al., 2018; Jardine and Litvak, 2003). However, 10 mM Mtz treatments extending beyond 24 hrs become increasingly toxic (Mathias et al., 2014) and removing Mtz from microtiter plates after a 24 hr pulse could not be easily automated. We therefore sought a 48 hr Mtz treatment regimen sufficient for inducing maximal rod cell loss by 7 dpf that showed no evidence of toxicity.

Five concentrations of Mtz were tested across a 2-fold dilution series from 10 mM to 625 μM. YFP reporter signals were quantified daily from 5-8 dpf using ARQiv. Changes in YFP levels were calculated as percentages normalized to non-ablated YFP controls. The data showed concentration-dependent reductions in YFP with maximal loss observed at 7 dpf for 10, 5, 2.5, 1.25 and 0.625 mM Mtz exposures; effect sizes of 55, 79, 87, 87, and 83 percent cell loss, respectively (Figure 1A; Supplemental Table 1). Although no lethality was observed for any condition, signs of distress were evident for 10 Mtz exposures (i.e., reduced motility). At ≤5 mM, however, no signs of stress were observed. As 2.5 mM Mtz was the lowest concentration producing maximal cell loss, this condition was selected as the treatment regimen for our large-scale screen.

**Figure 1.**
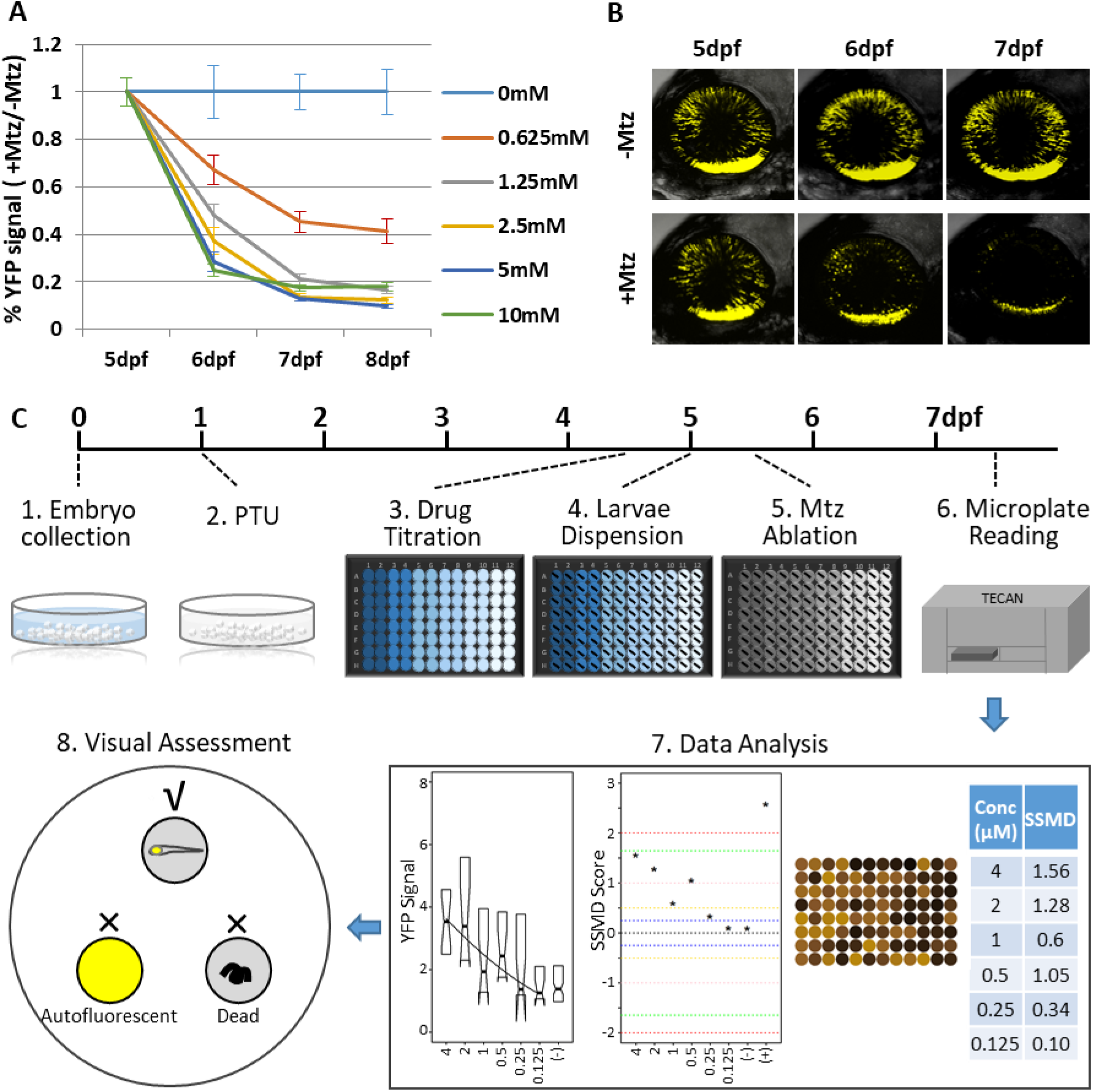
Zebrafish RP model and the schematic diagram of large-scale JHDL screening. (**A**) Optimization of Mtz treatment regimen for the inducible rod photoreceptor degeneration model of RP. Transgenic zebrafish larvae expressing a YFP-NTR fusion protein exclusively in rod photoreceptor cells were exposed to one of five concentrations of Mtz, from 10 mM to 0.625 mM, from 5 to 8 dpf (sample size: 56 larvae per group). YFP signal intensity for each larva was quantified daily using a microplate reader assay. The average YFP signal (±sem) of each Mtz treatment group is plotted as the percentage of non-ablated 0 mM Mtz controls per day. The 10, 5, and 2.5 mM Mtz-treated groups produced minimal YFP signal intensities (<20%) at 7 dpf (Supplementary Table 1). The 2.5 mM Mtz treatment condition was chosen for the primary screen. (**B**) Time series intravital confocal images of representative non-ablated (-Mtz control, upper panel) and 2.5 mM Mtz-treated (+Mtz control, lower panel) retinas from 5-7 dpf. By 7 dpf, only a limited number of YFP-positive cells are detectable in the +Mtz retina, mainly concentrated in a ventral band of high rod cell density. (**C**) Schematic of primary drug screening process: 1) At 0 dpf, large numbers of embryos were collected. 2) At 16 hpf, PTU was added to suppress melanization. 3) At 4 dpf, individual drugs were dispensed and titrated in 96-well plates using robotic liquid handlers; 16 wells (2 columns) per concentration and 6 concentrations total per drug. 4) At 5 dpf, COPAS was used to dispense individual larvae into single wells of 96-well plates. 5) After a 4 hr pre-exposure to drugs, larvae were treated with 2.5 mM Mtz. 6) At 7 dpf, YFP signals were quantified by microplate reader. 7) Same day data analysis using a custom R code to create a signal to background ratio plot, a SSMD score plot, a heat map of signal to background ratio, and a SSMD score table. 8) Drug plates producing a SSMD score ≥1 were visually inspected using fluorescence stereomicroscopy to exclude autofluorescent and lethal compounds.

Previously established power analysis methods using ablated and non-ablated controls (White et al., 2016) determined that a sample size of nine larvae was sufficient to detect a 50% neuroprotective effect. For ease of dispensing, microtiter plate formatting, and to account for larval dispensing errors, we increased the sample size to 16 larvae per condition for the primary screen. Fortuitously, this also allowed us to detect subtler neuroprotective effects. The strictly standardized mean difference quality control (SSMD QC) score was 1.67, indicating the assay was of sufficient quality to justify a large-scale screening effort (Zhang, 2011).

To establish a positive control, we tested 17 compounds and one compound “cocktail” previously implicated as retinal neuroprotectants (Supplementary Table 2). Unfortunately, none of these were able to sustain YFP expression at the concentrations tested (4 μM to 125 nM). However, a compound identified as a retinal cell neuroprotectant by the Zack lab (manuscript in preparation) did show-dose-dependent effects on YFP levels. This compound was therefore used as a positive control (POS) for assay performance.

### Primary Screen

For the large-scale PDD screen, the Johns Hopkins Drug Library (JHDL) was used. The JHDL is comprised of nearly 3,000 compounds, most being human-approved drugs (Shim and Liu, 2014). To minimize false discovery rates, all compounds were tested using qHTS principles (Inglese et al., 2006) – i.e., across six concentrations (4 μM - 125 nM) using a two-fold dilution series. The screen largely followed published ARQiv-HTS methodologies (G. Wang et al., 2015; White et al., 2016), with assay-specific details provided in Figure 1C (steps 1-8). In all, 2,934 compounds were screened and more than 350,000 transgenic zebrafish larvae evaluated. Real-time data analysis was performed as previously detailed (White et al., 2016) to generate: 1) a plot of YFP signal levels, 2) a plot of SSMD scores across all tested concentrations, 3) a signal intensity heat map of each plate, and 4) an SSMD score table (Figure 1C, step 7). Compounds producing SSMD scores of ≥1 were considered potential hits and flagged for visual inspection to assess fluorescence and general morphology using a stereo fluorescence microscope. This step facilitated elimination of false-positive compounds producing autofluorescence due to larval toxicity (5 drugs) or compound fluorescence (32 drugs) (Figure 1C, step 8). Additionally, this allowed visual confirmation of sustained YFP expression within the retina. At the conclusion of the primary screen, 113 compounds were identified as potential hits (Supplementary Table 3). Hits were classified according to the highest SSMD score achieved across all concentrations, and whether concentration-dependent effects were observed. SSMD scores suggested one drug produced a strong effect (SSMD of 2-3); four had semi-strong effects (1.645-2), 20 showed moderate effects (1.28-1.645), and 89 had semi-moderate effects (1-1.28) (Supplementary Table 3). Forty-two drugs showed concentration-dependent effects, while 72 exhibited discontinuous or singular concentration effects.

### Validation Assay I: Confirmation

We next performed a series of confirmatory and orthogonal assays to evaluate a subset of 42 hit compounds prioritized by SSMD score, dose-response profile, and/or implicated mechanism of action (MOA; Supplementary Table 4). Having extensive MOA data is a key advantage afforded by testing human-approved compounds which we leveraged in a previous large-scale zebrafish PDD screen (G. Wang et al., 2015). Similarly here, as studies have suggested inflammation plays a key role in retinal degeneration and regeneration (Hollyfield et al., 2008; Mitchell et al., 2018; White et al., 2017; Yoshida et al., 2013), hits implicated as modulators of inflammatory signaling were included. In addition, several compounds that did not produce concentration-dependent effects were selected to test whether this criterion was useful in predicting reproducibility. All compounds were obtained from new sources to ensure reagent authenticity. To confirm activity, three biological repeats were conducted but using a wider concentration range (from 100 μM to 1.28 nM using a 5-fold dilution series) to account for differences in reagent quality. If toxicity was observed at higher concentrations, dilution series were initiated at 10 µM or 1 µM. Using this strategy, 11 of the original 42 prioritized hit compounds were confirmed as lead compounds (26%; Figure 2). Effect sizes ranged from a 9 to 38 percent increase in YFP signal relative to +Mtz/0.1% DMSO controls (Fig. 2; drug abbreviations in Table 1).

**Figure 2.**
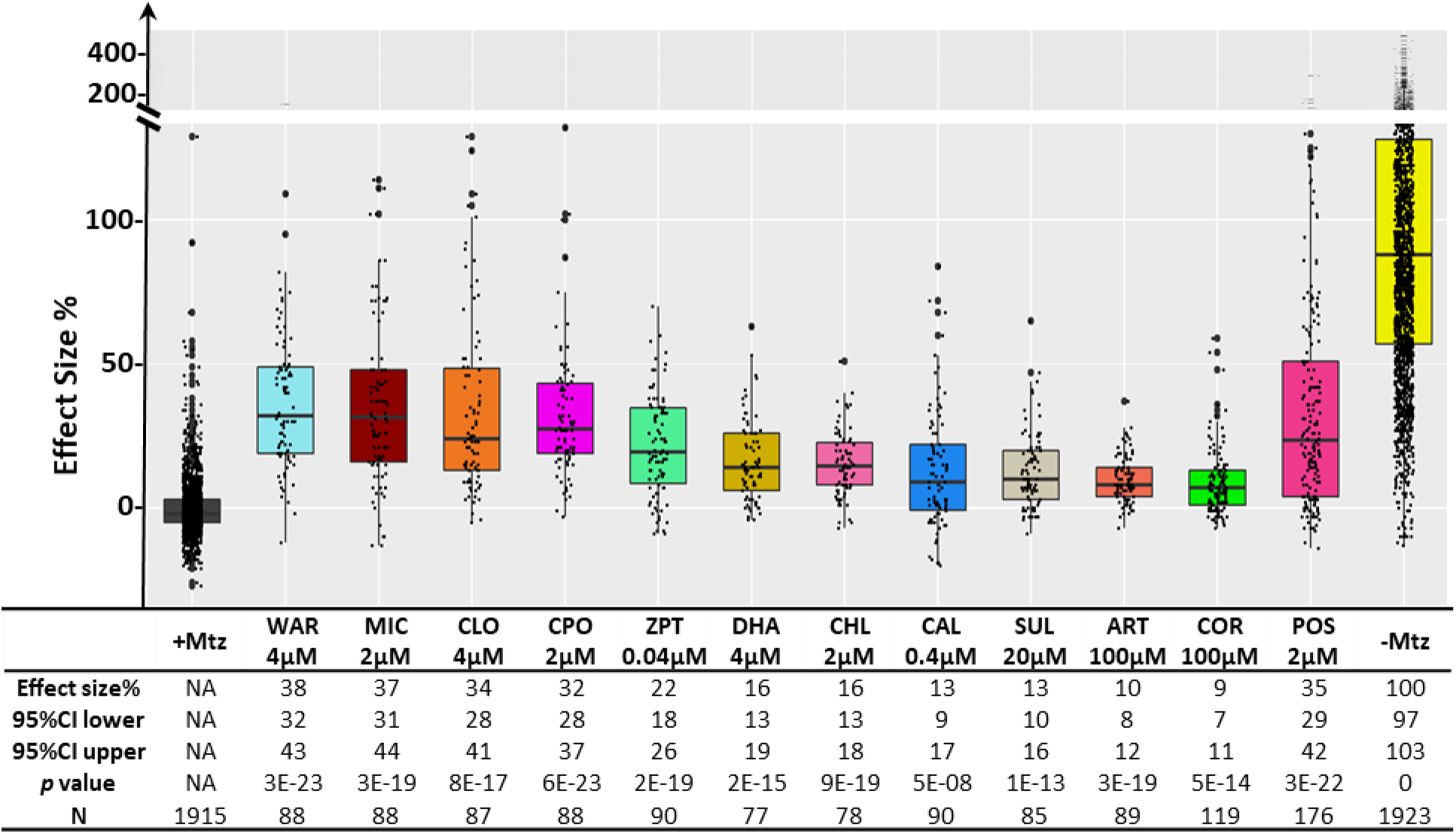
Neuroprotective effects of validated lead compounds. Box plots of eleven confirmed lead compounds and the positive control compound arrayed by neuroprotective effect size. YFP signal intensities ranged from a 9 to 38 percent increase over +Mtz controls. All experimental YFP values were normalized to the signal window calculated from the +Mtz controls (ablated, set at 0 percent) and -Mtz controls (non-ablated, set at 100 percent) to account for: 1) individual variation, 2) fluctuations in signal window per assay, and 3) to allow data from identical conditions to be pooled across assays; effect sizes, 95% confidence intervals, *p* values, and sample sizes (N) for each condition are shown below. *P* values were calculated by comparing drug-treated conditions to +Mtz controls using Student’s *t* test followed by Bonferroni correction for multiple comparisons (α=0.004 adjusted significance level). No statistical differences in larval survival were observed for lead compounds relative to their respective +Mtz controls, except for DHA (86%) and CHL (87%; Fisher’s exact test, p<0.05).

**Table 1.**
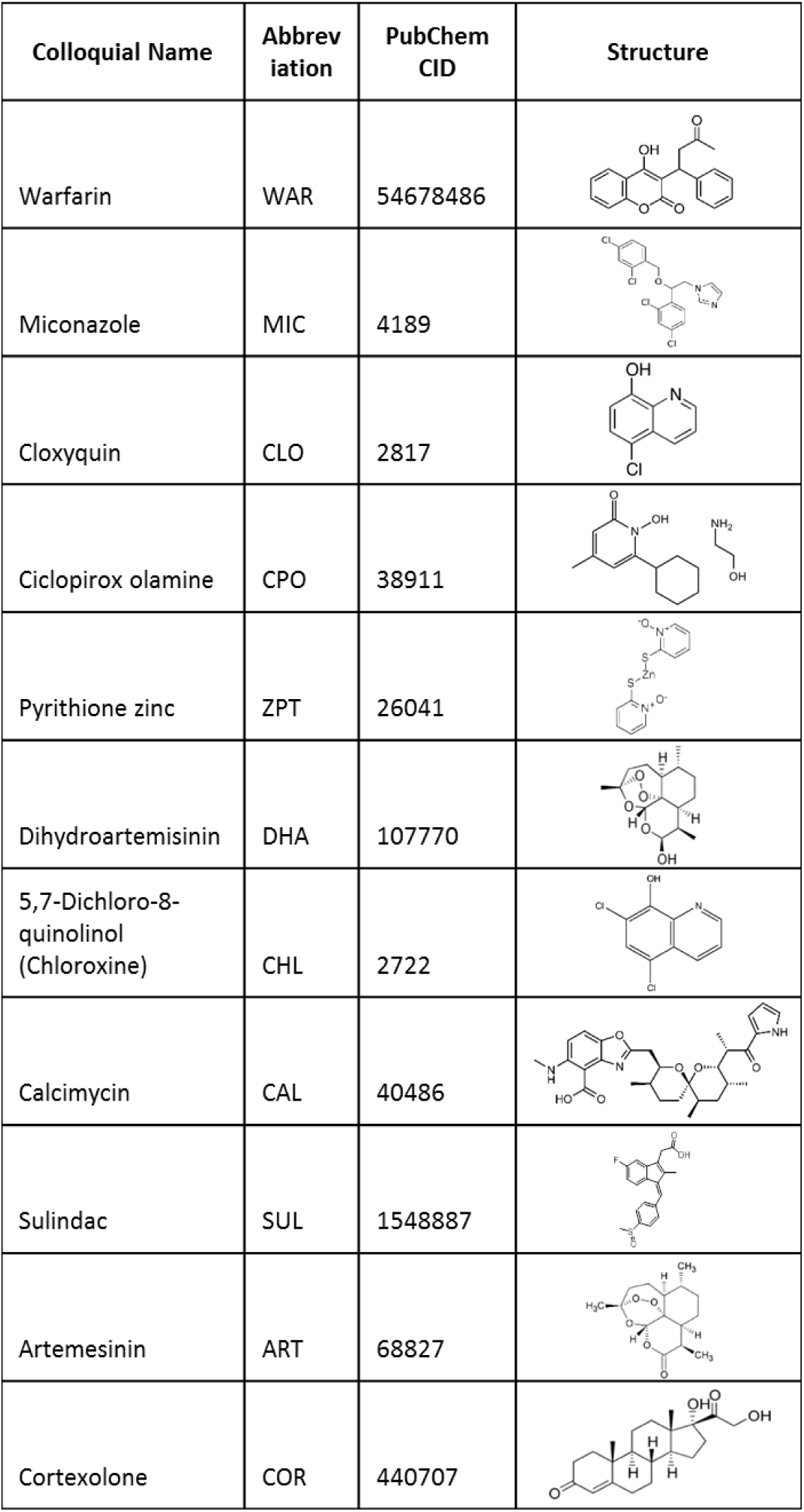
Eleven lead compounds. . Abbreviations, PubChem CID, and chemical structures of the validated neuroprotective compounds.

We next asked whether there was a correlation between SSMD scores and/or concentration-dependent effects and confirmation rates. Among 19 selected compounds with higher SSMD scores (≥1.3), seven (37%) were confirmed; among 23 with lower SSMD scores (1-1.28), four (17%) were confirmed. Of 27 compounds with a concentration-dependent trend, eight (30%) were confirmed. Conversely, of 15 compounds that did not show a concentration-dependent trend, 3 (20%) were confirmed. Among the eleven confirmed leads, eight (73%) showed dose-dependent effects and seven (64%) had higher SSMD scores (>1.28). These results suggest that prioritizing hit compounds by both relative SSMD score and dose-dependent trends provides predictive value for confirming activity, consistent with qHTS principles (Inglese et al., 2006).

### Validation Assay II: Confocal intravital microscopy

To ensure lead compounds promoted rod photoreceptor cell survival rather than simply increased YFP signal intensity, intravital time series confocal microscopy was used to image lead-treated, +Mtz and -Mtz control retinas. Mtz-ablated retinas exhibited dramatically reduced YFP signal (Figure 3, +Mtz). Conversely, rod photoreceptor cells in non-ablated -Mtz control retinas displayed robust YFP signal throughout the retina and elongated morphologies suggesting healthy outer segments (Figure 3, -Mtz). In retinas exposed to Mtz and lead compounds, YFP signal loss was attenuated and rods typically displayed healthy elongated morphologies (Figure 3, e.g., CLO). However, some cells appeared rounded, suggesting that the process of degeneration was not fully inhibited, e.g., miconazole (MIC; Figure 3). To confirm increased rod cell survival in lead drug treated groups (versus increased YFP intensity), confocal images of YFP-expressing cells were 3D-rendered and fluorescence volumetrically quantified using Imaris software-based automated image analysis (White et al., 2017). The data showed YFP volumes were elevated in all drug treated groups relative to Mtz-ablated controls (Supplementary Figure 2), confirming that lead compounds promoted increased rod cell numbers and/or preserved outer segment morphology.

**Figure 3.**
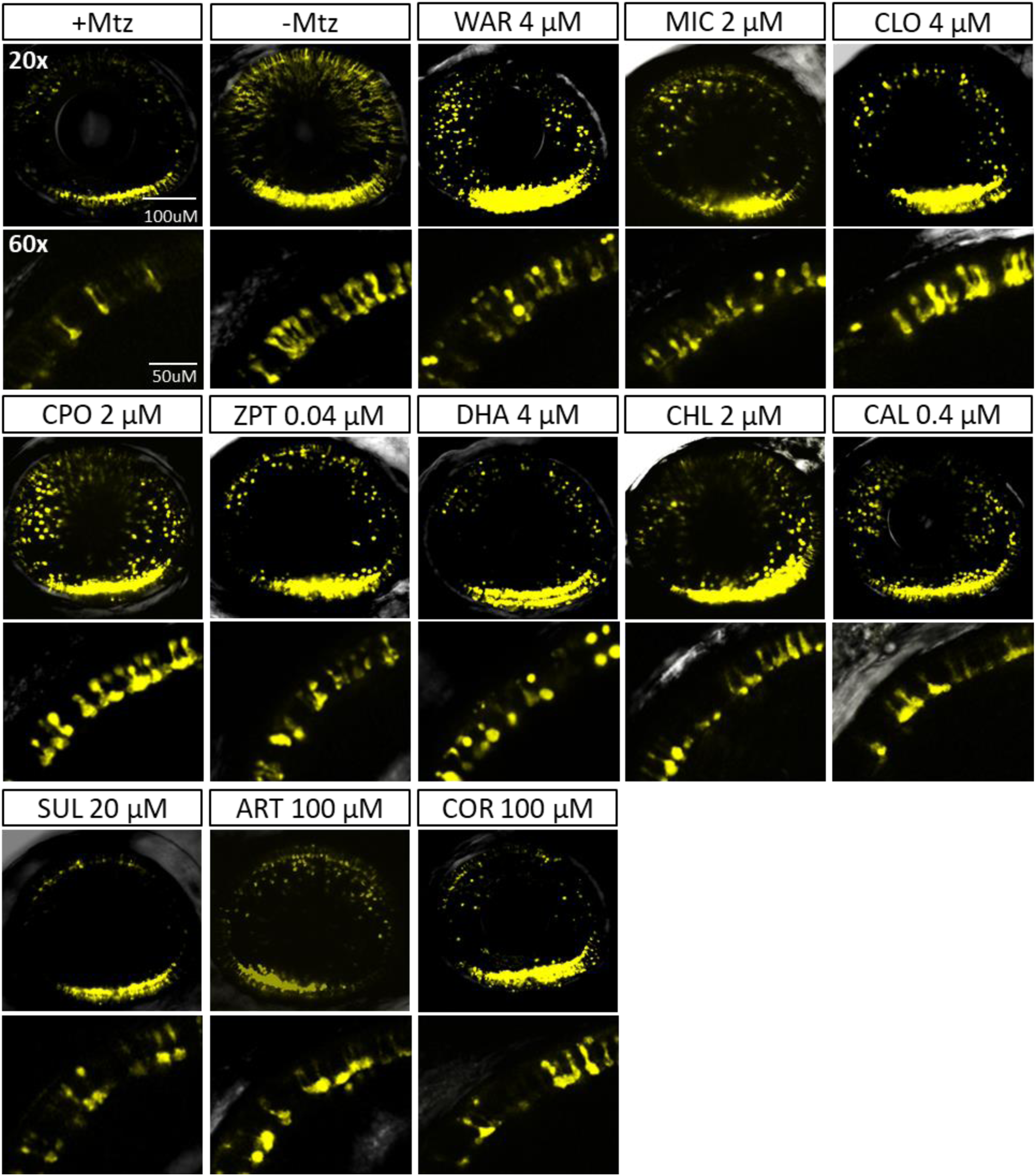
Confocal images of neuroprotective effects of confirmed lead drug compounds. Representative intravital whole retina confocal image stacks of +Mtz (ablated control), -Mtz (non-ablated control), and lead compound-treated retinas taken with a 20× (upper panels) and 60× objective (lower panels) at 7 dpf. The 20× objective images show loss or preservation of YFP signals throughout the retina, while the 60× objective images provide morphological detail of YFP-expressing rod cells. The +Mtz control shows extensive YFP signal loss, with remaining cells restricted largely to the ventral region. The -Mtz control shows unperturbed YFP signal throughout the retina. Drug-treated retinas exhibit sustained YFP signals and preservation of rod photoreceptor morphology to varying extents.

### Validation Assay III: NTR inhibition

As rod cell death is induced by NTR reduction of the prodrug Mtz in our model, it is possible that some lead compounds simply suppressed NTR enzymatic activity. To test this, NTR activity was evaluated in the presence of each lead compound by assaying the reduction kinetics of the prodrug CB1954 *in vitro* (Prosser et al., 2010). To ensure any potential for NTR inhibition was accounted for, all compounds were tested at 300 μM (∼100-fold greater than neuroprotective concentrations). Compounds were deemed potential inhibitors if NTR activity was less than 75% of the control. Seven compounds showed no evidence of NTR inhibition by this criterion, but four did: warfarin (WAR), ciclopirox olamine (CPO), calcimycin (CAL) and sulindac (SUL) (Supplementary Figure 3). However, IC_50_ measures ranged from 150 µM (for CPO) to 350 µM (for SUL), substantially higher than neuroprotective concentrations (i.e., 0.4-20 µM, see Figure 2).

When NTR reduction of Mtz was tested, similarly weak inhibitory effects were observed (Supplementary Figure 3B). The differences in concentrations between neuroprotective and NTR inhibitory activities diminish the possibility that leads act directly on NTR. To test of this further, lead compounds were assayed for neuroprotective effects in NTR-independent mouse RP models assays (see below).

### Validation Assay IV: Rod photoreceptor development

To control for the possibility that confirmed compounds promoted rod photoreceptor development, rather than provided neuroprotection, YFP levels were quantified in *rho:YFP-NTR* larvae exposed solely to lead drugs from 5-7 dpf. Retinoic acid (RA, 1.25 μM) was used as a positive control as it promotes rod fates during development in zebrafish (Hyatt et al., 1996). RA treated fish displayed significantly increased YFP signals compared to untreated controls (Supplementary Figure 4). In contrast, none of the retinas treated with lead compounds exhibited increased YFP expression, suggesting they do not promote rod photoreceptor cell fate. Interestingly, three lead compounds cloxyquin (CLO), cortexolone (COR) and CPO produced lower YFP signals than controls, suggesting negative effects on rod cell development.

### Validation Assay V: Regeneration

It is well known that the zebrafish retina regenerates (Gorsuch and Hyde, 2014; Lenkowski and Raymond, 2014; Wan and Goldman, 2016). Therefore, to determine whether lead compounds acted by stimulating regeneration, we used a previously described ARQiv assay designed to detect changes in rod cell replacement kinetics (Walker et al., 2012; White et al., 2017). Briefly, *rho:YFP-NTR* larvae were first treated with 10 mM Mtz at 5 dpf for 24 hrs to induce rod cell loss. At 6 dpf, Mtz was washed out and larvae were treated with lead compounds at concentrations corresponding to maximal neuroprotective effects and YFP levels quantified at 9 dpf. Dexamethasone, which accelerates rod cell regeneration kinetics (White et al., 2017), was used as a positive control. The results showed that none of the compounds increased rod cell regeneration rates (Supplementary Figure 5). Interestingly, four compounds, dihydroartemisinin (DHA), CLO, CPO and MIC inhibited regeneration. These data suggest that lead compounds do not increase YFP levels by promoting rod cell regeneration.

### Molecular Mechanism of Action

Previously, we used only “on label” information to explore MOA of hit compounds (G. Wang et al., 2015); e.g., Supplementary Table 4). Recently, we have become interested in additional advantages afforded by whole-organism PDD, such as polypharmacology (Dar et al., 2012; Rennekamp and Peterson, 2015; Rihel et al., 2010). Accordingly, here we also applied a target-agnostic MOA analysis process by evaluating lead compound performance in HTS/uHTS studies archived on PubChem (https://pubchem.ncbi.nlm.nih.gov/). Many lead compounds exhibited shared target activities, suggesting common MOA (Supplementary Table 5). The most common target implicated was Tyrosyl-DNA Phosphodiesterase 1 (TDP1), a DNA repair enzyme. To test whether TDP1 inhibition was a viable means of promoting rod photoreceptor survival, we tested three TDP1 inhibitors in our zebrafish RP model: paromycin (PAR), thiostepton (THI) and methyl-3,4-ephostain (MET) (Huang et al., 2011; Liao et al., 2006). Both PAR and THI showed neuroprotective effects (Table 2, Figure 4).

**Figure 4.**
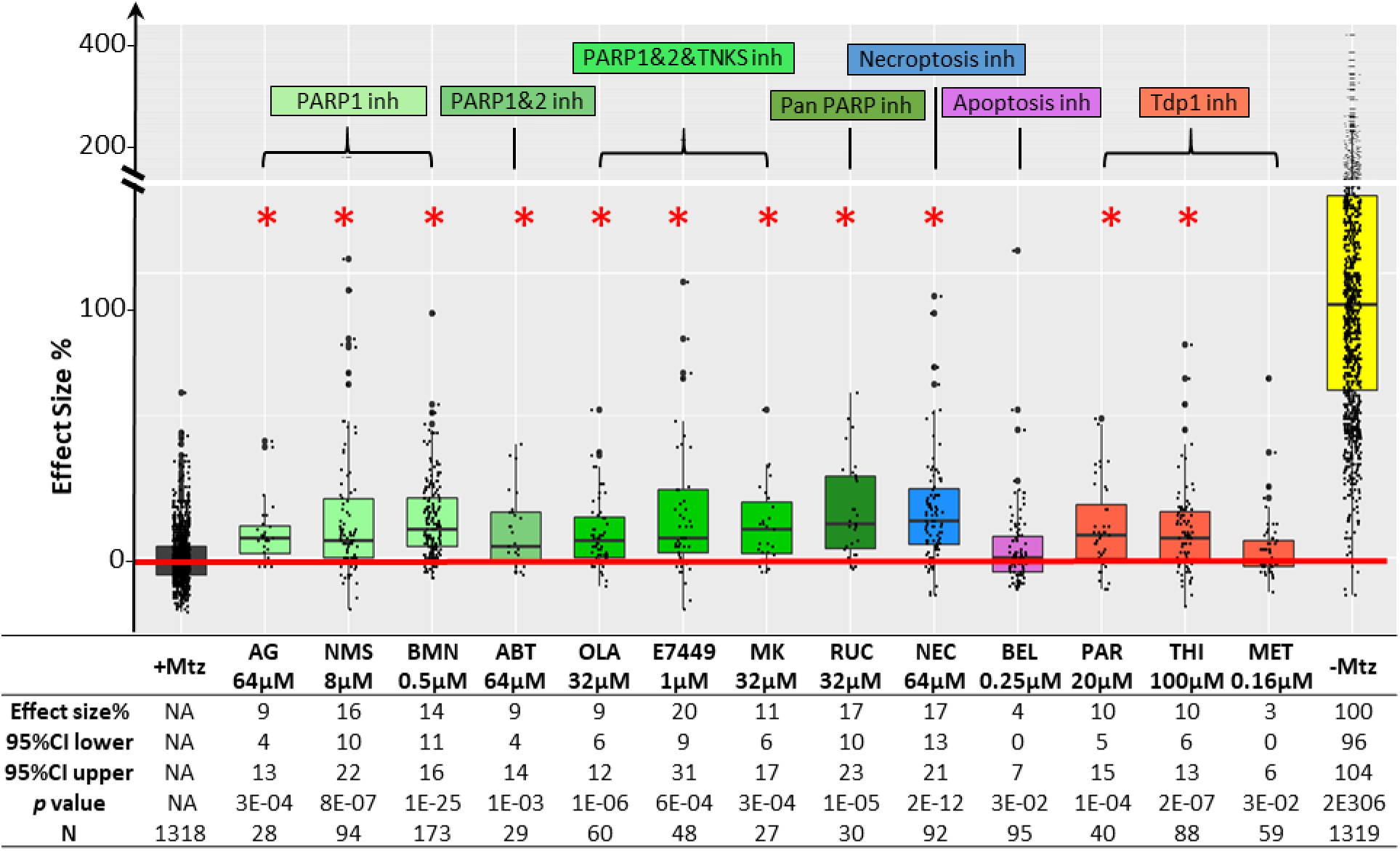
Analysis of potential mechanisms of action (MOA). Box plots of PARP inhibitors (green), necroptosis inhibitor (blue), apoptosis inhibitor (magenta), and TDP1 inhibitors (orange) tested for neuroprotective effects in the zebrafish RP model as per the primary screen. All PARP inhibitors, two of three TDP1 inhibitors, and the necroptosis inhibitor promoted rod cell survival. The individual effect sizes, 95% confidence intervals, *p-*values, and sample sizes (N) for each group are shown below. *P* values were calculated by comparing drug-treated conditions to +Mtz controls using Student’s *t* test followed by Bonferroni correction for multiple comparisons (α=0.003 adjusted significance level). AG: AG-14361; NMS: NMS-P118; BMN: Talazoparib; ABT: Veliparib; OLA: Olaparib; MK: Niraparib; RUC: Rucaparib; NEC:Necrostatin-1; BEL: Belnacasan; PAR: Paromomycin; THI: Thiostrepton; MET:Methyl-3,4-ephostatin. No statistical differences in larval survival were observed for tested compounds relative to their respective +Mtz controls, except for PAR (67%; Fisher’s exact test, p<0.05).

**Table 2.**
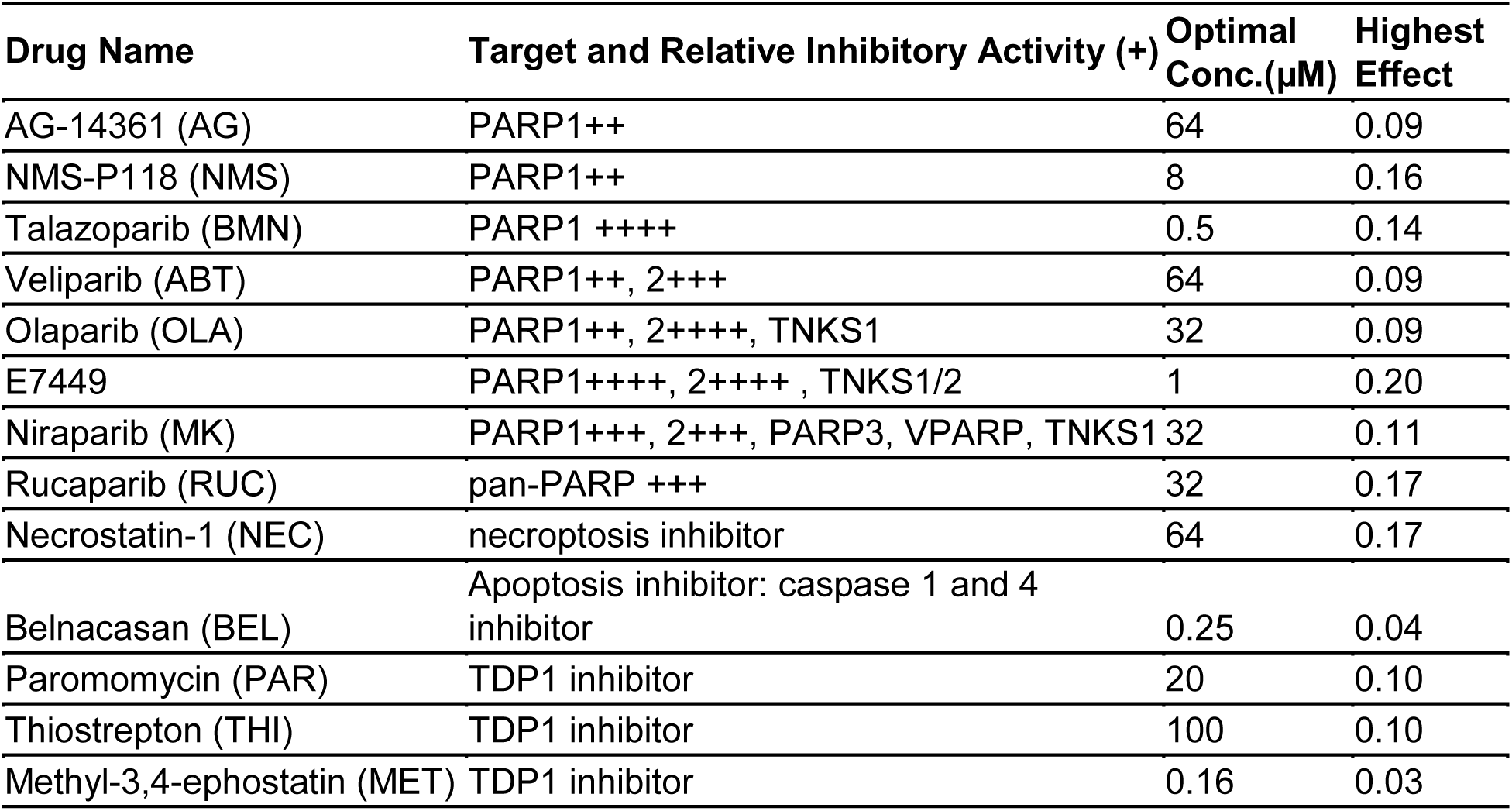
HTS-implicated targets and cell death pathway inhibitors tested. . Eight PARP inhibitors, one necroptosis inhibitor, one apoptosis inhibitor and three TDP1 inhibitors were tested for neuroprotective effects in zebrafish. Target specificity and relative inhibitory activity of PARP inhibitors are as reported by Selleckchem (https://www.selleckchem.com/PARP.html). The concentration producing the highest effect size and the maximal rod cell survival rate achieved are listed for each compound. Abbreviations: TDP1 (Tyrosyl-DNA phosphodiesterase 1), PARP (Poly (ADP)-ribose Polymerase), TNKS (Tankyrase), VPARP (vault PARP, aka PARP4). Note: TNKS1 and TNKS2 are also known as PARP5 and PARP6, respectively.

This result is surprising given that NTR reduction of Mtz is thought to cause DNA damage-induced cell death (Curado et al., 2007). Thus, inhibition of a DNA repair enzyme would be expected to enhance NTR/Mtz-mediated cell death not inhibit it. However, an alternative means of disrupting TDP1 activity, is by inhibiting Poly (ADP-ribose) Polymerases (PARPs; Murai et al., 2014). PARPs also mediate DNA repair but, interestingly, hyperactivation of PARP1 leads to a specific form of cell death, termed parthanatos (Fatokun et al., 2014; Wang et al., 2016). We therefore assayed PARP inhibitors for the capacity to promote rod cell survival. All eight PARP inhibitors tested had neuroprotective activity, ranging from 9 to 20% (Figure 4). To account for other cell death mechanisms implicated in neurodegeneration, we tested inhibitors of necroptosis (necrostatin-1; NEC), and apoptosis (belnacasan; BEL). Surprisingly, NEC promoted rod cell survival, while BEL did not (Figure 4, Table 2).

### Mouse Model Validation I: primary retinal cells treated with stressor compounds

Our ultimate goal is to identify potential new therapeutics for RP patients. We reasoned compounds that show similar effects in fish and mammalian models would be more likely to translate to human RP patients. Therefore, we tested the efficacy of lead compounds in mouse models of retinal degeneration.

First, we tested compounds for the capacity to protect primary mouse retinal cells from stress-induced cell death in culture. Retinal cells were isolated from postnatal day four (P4) wildtype mice and grown as previously described (Fuller et al., 2014). To induce retinal cell death, tunicamycin (0.6 µg/mL) or thapsigargin (0.25µM) were added to the cultures. These “stressor” compounds inhibit protein glycosylation (Fliesler et al., 1984) and endoplasmic reticulum (ER) calcium levels (Thastrup et al., 1990), respectively (E. Lai et al., 2007). In turn, they induce an unfolded protein response (UPR) (H. Wang et al., 2015) and related ER stress (Oslowski and Urano, 2011; Zhang et al., 2014) which have been implicated in the etiology of RP (Griciuc et al., 2011)(Rana et al., 2014). For pan-retinal cell survival assays, compounds were tested at 5 concentrations across a 2-fold dilution series (from 4 µM to 250 nM) in sextuplicate. After two days in culture, relative viability was quantified using a luminescent cell viability assay (CellTiter-Glo). A second assay, for photoreceptor survival effects was performed using cells isolated from QRX mice (Wang et al., 2004), a transgenic line in which GFP expression is restricted to photoreceptors. For these assays, compounds were tested at 7 concentrations across a 3-fold dilution series (from 30 µM to 40 nM) in sextuplicate. After 48 hrs, cells were imaged and analyzed using a high-content screening system (Cellomics VTI). The photoreceptor survival was assessed by quantifying the number of GFP-expressing cells. For both assays, compounds were considered effective if the mean number of surviving total retinal cells or photoreceptor cells was two standard deviations greater than the mean of the non-drug treated control. Six hit compounds, CAL, CPO, DHA, WAR, chloroxine (CHL) and pyrithione zinc (ZPT) showed protective effects in at least one of the two assays (Table 3). CPO and ZPT protected photoreceptors from both tunicamycin and thapsigargin-induced stress (Table 3).

**Table 3.** Mouse primary retinal cell culture tests. . Isolated primary retinal cells were cultured for 2 days in the presence of tunicamycin (0.6 µg/ml) or thapsigargin (0.25 µM) to induce ER stress, after which cell survival was measured. (A) Three compounds promoted retinal neuron survival in non-sorted cultures (denoted by *). (B) Four compounds promoted the survival of GFP-labeled photoreceptor cell specifcally (denoted by *). WAR: warfarin, CPO: ciclopirox olamine, ZPT: pyrithione zinc, DHA: dihydroartemisinin, CHL: chloroxine, CAL: calcimycin, ART: artemisinin, COR: cortexolone, SUL: sulindiac. (++: >2SD; +++: >3SD)

| A | Drug Name | Retinal cell<br>(general cell loss) |  |
| --- | --- | --- | --- |
|  |  | Tunicamycin | Thapsigargin |
| * | WAR | ++ | ++ |
| * | CPO | - | +++ |
| * | DHA | - | ++ |
|  | SUL | - | - |

**Table 3. Mouse primary retinal cell culture tests.**
| B | Drug Name | Photoreceptor-GFP<br>(cell specific loss) |  |
| --- | --- | --- | --- |
|  |  | Tunicamycin | Thapsigargin |
|  | WAR | - | - |
| * | CPO | ++ | ++ |
| * | ZPT | ++ | ++ |
|  | DHA | - | - |
| * | CHL | - | ++ |
| * | CAL | - | ++ |
|  | ART | - | - |
|  | COR | - | - |

### Mouse Validation II: rd1 mutant retinal explants

We next examined the effects of lead compounds in mouse retinal explants isolated from *rd1* mice. The *rd1* mouse model of RP exhibits early onset rod cell degeneration caused by a mutation in the *Pde6b* gene (Chang et al., 2002), an ortholog of human RP-associated gene *PDE6B* (Bayés et al., 1995; Gal et al., 1994; Hmani-Aifa et al., 2009; McLaughlin et al., 1993). In these mice, photoreceptor degeneration begins around P10 and progresses rapidly. By P21, only a few rows of photoreceptor cells remain in the outer nuclear layer (ONL; LaVail and Sidman, 1974). Here, retinal explants from P10 *rd1* mice were isolated and cultured *ex vivo* (Bandyopadhyay and Rohrer, 2010). Eight lead compounds were evaluated at three concentrations across a five-fold dilution series centered on the most effective concentration determined in fish RP models. After eleven days in culture, explants were fixed, the retinal cells stained and the number of photoreceptor rows counted. Neuroprotective effects were defined as exhibiting 1) a concentration-dependent increase in the number of photoreceptor rows remaining in the ONL relative to untreated controls and 2) a *p*-value of ≤0.05. An average of 1.2 ±0.19 rows of photoreceptors remained in the ONL of non-treated control explants cultured for eleven days. Three of eight tested drugs, CPO, DHA and artemisinin (ART), increased the number of surviving photoreceptor layers, suggesting cross-species conservation of neuroprotective effects (Figure 5). However, high concentration CPO treatments (15 µM) led to disruption of retinal histology due to induction of proliferation in the inner nuclear layer (INL) and ONL.

**Figure 5.**
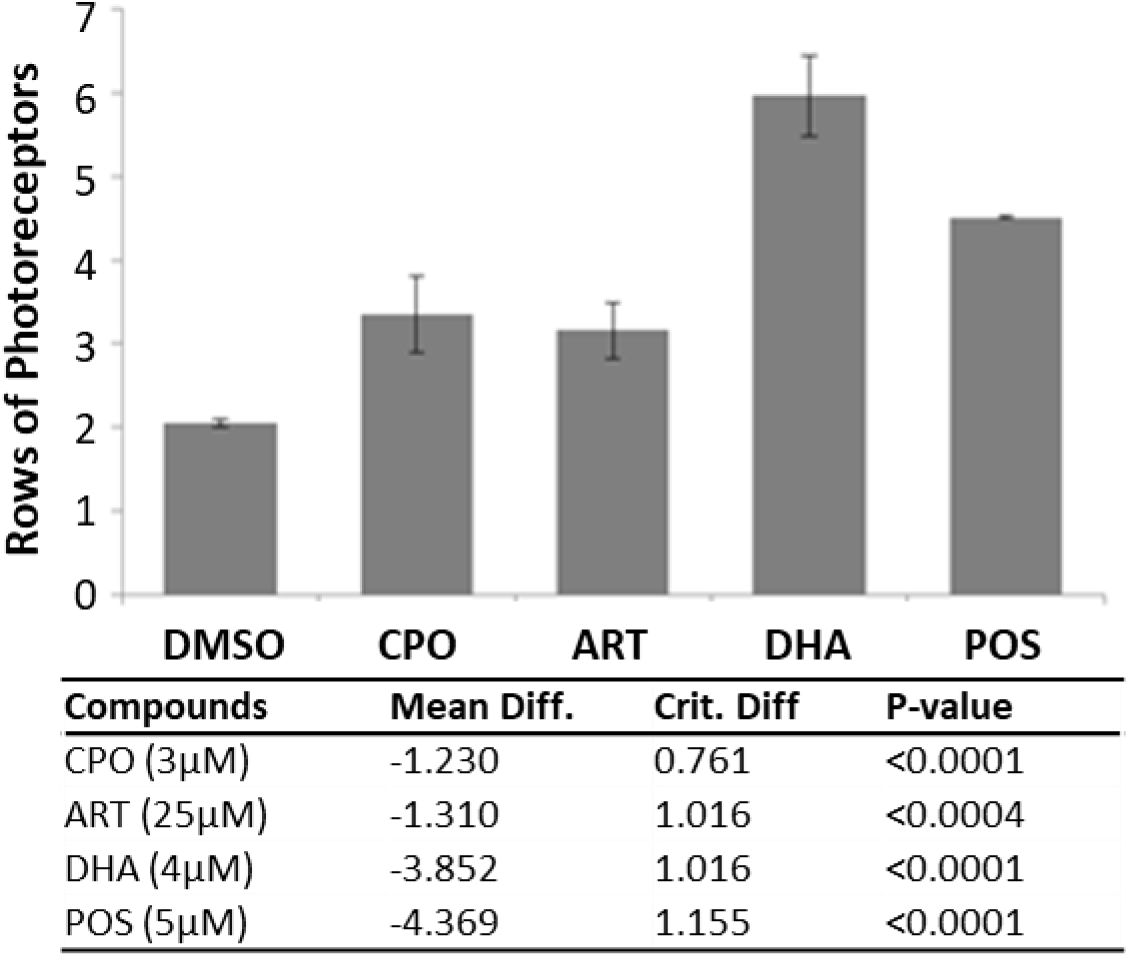
Tests of neuroprotective effects in mouse *rd1* retinal explants. Mouse r*d1* retinal explants were cultured for eleven days in the presence of lead drugs or DMSO, then fixed, sectioned, nuclear stained and the number of rows photoreceptors quantified. All compounds were screened at three different concentrations to test for dose-dependent effects (sample size of 3 to 4 explants per condition). Among 6 tested compounds, artemisinin (ART), dihydroartemisinin (DHA), and ciclopirox olamine (CPO) showed dose-dependent photoreceptor layer preservation. ANOVA followed by Bonferroni/Dunn test was performed to calculate *p* values comparing drug-treated samples to DMSO controls (*p*<0.005).

### Mouse Validation III: rd10 mutant model of RP

Finally, we examined the effects of the lead drug candidate DHA for neuroprotective effects in the *rd10* mouse model of RP (*Pde6b^rd10^*). The *rd10* line was selected for *in vivo* experiments due to its slower rate of photoreceptor degeneration relative to other *rd* mutants, thus allowing for a longer duration of time for pharmacological intervention. DHA was selected based on its superior performance in *rd1* retinal explant assays (Figure 6) and because it is the active metabolite of a second lead drug candidate, ART, an antimalarial compound. Prior to testing, DHA was encapsulated in PLGA polymers. Release kinetics assays in PBS/0.1% DMSO *in vitro* suggested DHA would reach maximal concentrations after 30 days, and remain stable for at least 20 days thereafter (Supplementary Figure 6). PLGA-DHA was injected into the vitreous of one eye of *rd10* mice at P14, with the contralateral eye serving as a vehicle injection control. At P32, eighteen days after injection, and when DHA levels were predicted to reach 80% of maximal concentration, *rd10* mouse eyes were processed for immunohistochemistry to quantify ONL thickness. Despite initial promising results in pilot assays, no reproducible neuroprotective effects were observed in these assays (Supplementary Figure. 7).

**Figure 6.**
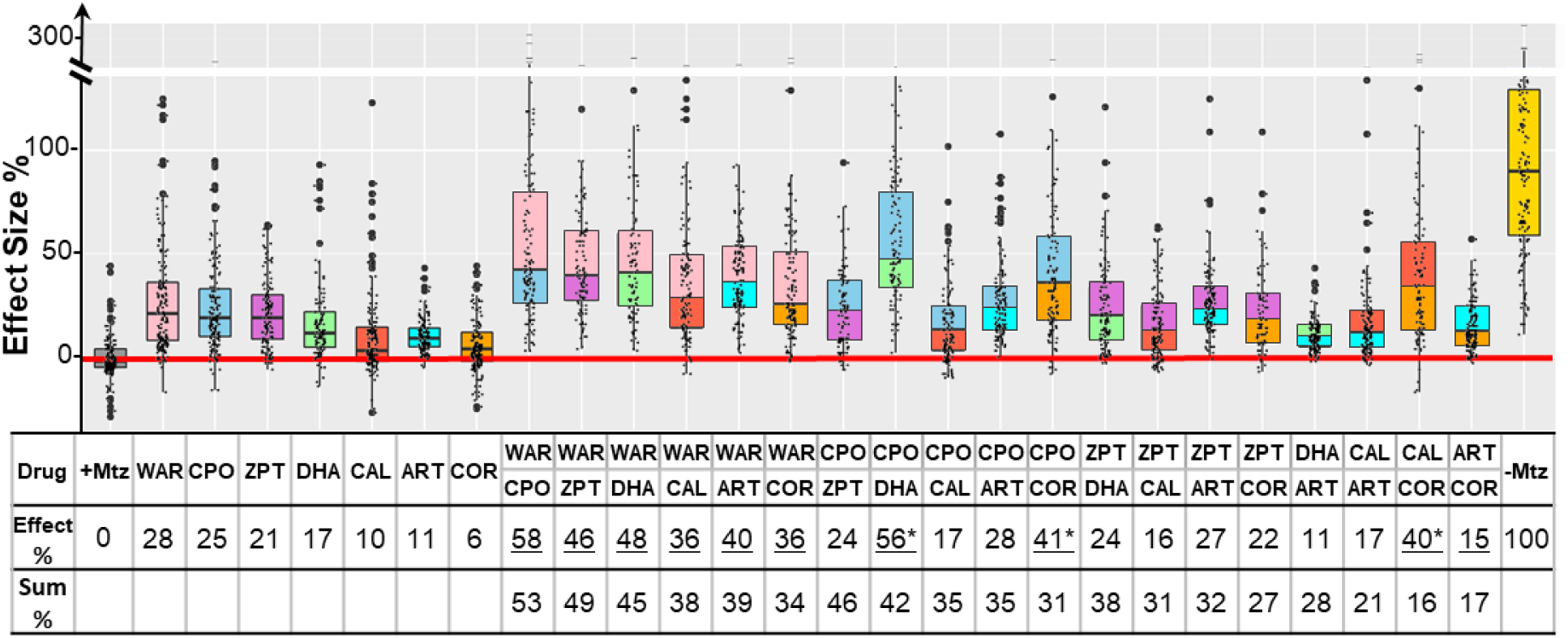
Additive effects of paired lead compounds. Seven lead compounds were screened in pairs for additive neuroprotective effects (i.e., combining individual maximal effective dosages and testing for enhanced survival effects). An additive effect was defined as any combinatorial treatment equal to or greater than the sum of the corresponding individual compound effects ±10%. Of twenty-one pairs tested, ten produced additive effects on rod cell survival (underscored). Maximal neuroprotective effects reached ∼58% survival, and three pairs produced better than additive effects (denoted by *). Note: two pairs, DHA+CAL and DHA+COR resulted in lethality and are therefore not plotted. Effect %: effect size of each paired treatment; Sum %: mathematically summed effect of the two individual treatments. Note, 95% confidence intervals, *p*-values, and sample sizes are provided in Supplementary Table 6. No statistical differences in larval survival were observed for lead compounds or compound pairs relative to their respective +Mtz controls, except for DHA (73%), WAR+DHA (73%), CIC+DHA (89%), ZPT+DHA (91%), ZPT+COR (90%) and CAL+COR (79%; Fisher’s exact test, p<0.05).

### Combinatorial Assay

PubChem searches also suggested potential complementary MOA, i.e., multiple independent targets across compounds (Supplementary Table 5). We therefore hypothesized that combining lead compounds may produce additive effects. To test this idea, seven lead compounds were tested in pairs using optimal effective concentrations (a total of 19 pairs; two being lethal). Pairs producing an effect equal to (±10%) or greater than the sum of their individual values were considered additive. By this criterion, 10 of 19 pairs exhibited additive effects (Figure 7, Supplementary Table 6).

**Figure 7.**
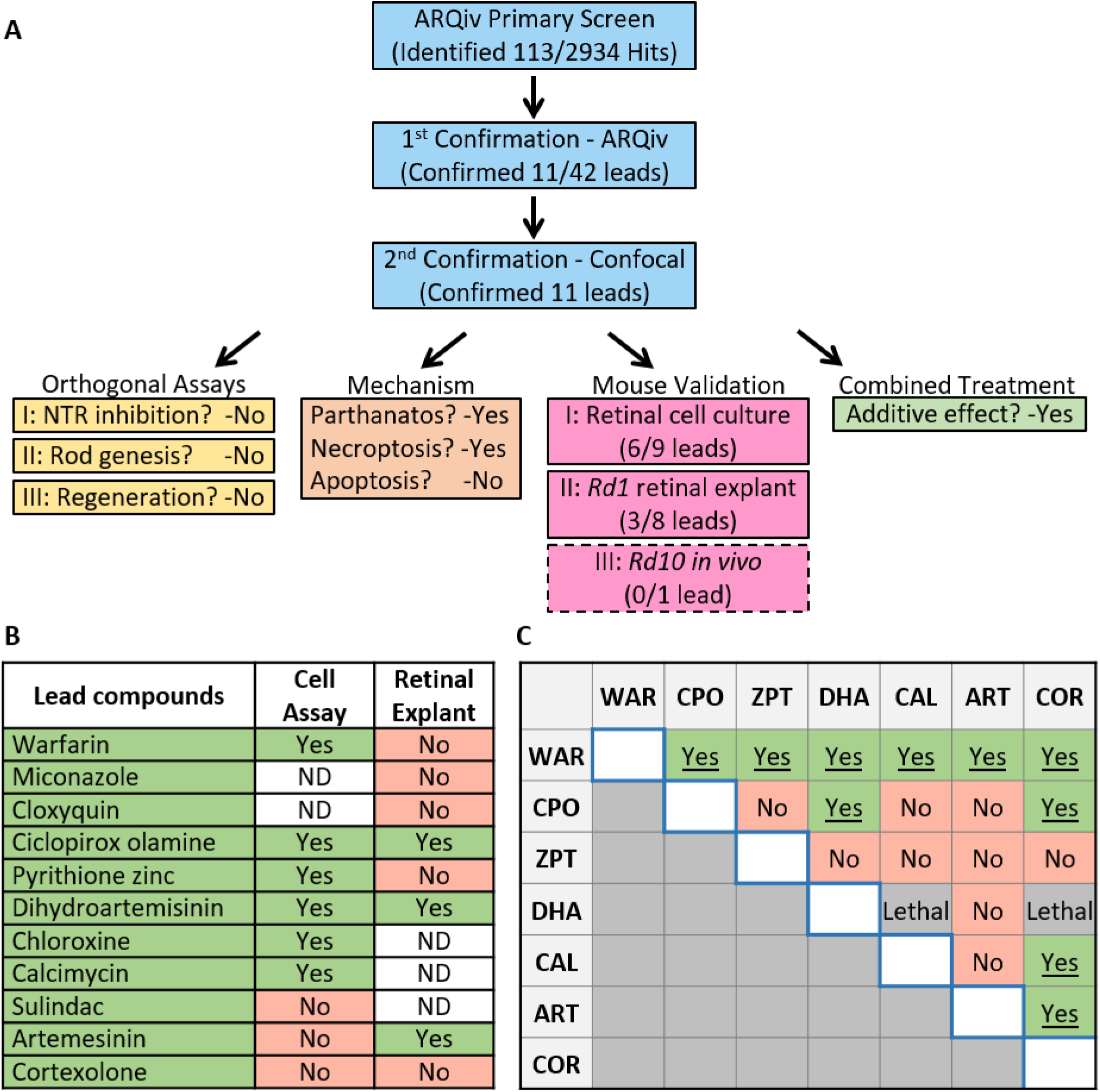
Summary. (A) Flow chart of PDD process: primary drug screen, secondary confirmations, orthogonal assays, MOA assays, cross-species testing and combinatorial paired drug tests. The primary screen implicated 113 “hit” compounds as neuroprotectants out of 2,934 tested; 11 out of 42 prioritized hits were confirmed as leads; 6 of 9 leads tested were validated in mouse primary retinal cell cultures, 3 of 8 were validated using *rd1* retinal explants, and two leads were effective in both culture systems (B); one lead failed when tested in *rd10* mouse retinas. Shared MOA analyses implicated PARP1-dependent cell death pathways, i.e., parthanatos and/or cGMP-dependent, as a common target. Combinatorial assays demonstrated additive effects (C), suggesting complementary neuroprotective MOA as well. Yes: effective; No: fail; ND: not determined.

Moreover, three pairs produced supra-additive effects (i.e., ≥25% greater than the sum) suggesting possible synergy (Figure 6, denoted by *). The maximum paired effect reached 58% rod cell survival. Several compounds showed broadly additive effects, e.g., WAR was additive with all six drugs and COR with four of five pairs (one pair proving lethal). Additive effects suggest multiple signaling pathways are involved in NTR/Mtz-induced photoreceptor degeneration. A schematic of the entire screening cascade is provided in Figure 7.

## DISCUSSION

Identifying effective neuroprotective therapies for RP and other IRDs stands as a critical unmet need for the field (Duncan et al., 2018; Wubben et al., 2019). Although, neurotrophic factors, anti-apoptotic agents, nutritional supplements and antioxidants have shown neuroprotective effects in animal models of RP (Dias et al., 2017).

Unfortunately, these reagents have produced, at best, only limited benefits for patients to date and, for some, mild improvements are offset by adverse side effects associated with long-term use (Dias et al., 2017). For example, ciliary neurotrophic factor (CNTF) was shown to be effective in protecting photoreceptors in mouse (Cayouette et al., 1998), dog (Tao et al., 2002) and chicken (Fuhrmann et al., 2003) models of retinal degeneration. However, CNTF failed to improve either visual acuity or field sensitivity in short- and long-term RP clinical trials (Birch et al., 2016, 2013; Ho et al., 2015). Clinical trials of Vitamin A in combination with Vitamin E (Berson et al., 1993), docosahexaenoic acid (Berson et al., 2004), lutein (Berson et al., 2010) or valproic acid (Birch et al., 2018) were reported to produce some benefits for RP patients, but only in subpopulations, and some of these studies have been controversial (Massof and Finkelstein, 1993).

Our strategy for addressing this challenge has two key elements: 1) scaling up the number of compounds tested directly in complex living disease models, and 2) a cross-species screening cascade that starts with small models amenable to HTS and proceeds to mammalian models. We hypothesize that compounds producing beneficial outcomes across evolutionarily diverse species will target conserved MOA and thus stand a higher chance of successfully translating clinically. As a generalized strategy, large-scale drug discovery screens using small animal models are showing increasing promise across multiple disease paradigms (Cagan et al., 2019; Cully, 2019; Kitcher et al., 2019; MacRae and Peterson, 2015).

Here, using a large-scale *in vivo* drug screening platform (Figure 1), we tested 2,934 largely human-approved compounds for neuroprotective effects across six concentrations in >350,000 larval zebrafish models of RP. The primary screen implicated 113 compounds as neuroprotectants (Supplementary Table 3). Confirmatory repeats and a series of four orthogonal assays validated 11 of 42 prioritized hit compounds in protecting zebrafish rod photoreceptors from cell death (Figure 2, 3, supplementary Figures 3, 4, 5 and Table 1). Importantly, investigations of lead compound MOA, led to the discovery that NTR/Mtz-mediated rod cell death appears to proceed through alternative cell death pathways recently linked to photoreceptor degeneration (Figure 4).

To further test relevance to disease mechanisms, and conservation of neuroprotective effects across species, lead compounds were evaluated in three mouse IRD/RP models. Six of nine leads were confirmed as neuroprotectants in primary retinal cell cultures (Table 3). Three of eight leads assayed using *rd1* retinal explant cultures (Figure 5); two being active in both paradigms, dihydroartemisinin (DHA) and ciclopirox olamine. We chose the *rd10* RP model for *in vivo* testing because it undergoes a slower rate of rod cell loss than *rd1*; spanning from approximately P16 to P35 (Chang et al., 2007; Gargini et al., 2007). DHA was the most promising compound for these tests as it had shown strong effects across assays and was amenable to a long-term release formulation designed to sustain drug action over weeks to months (Supplementary Figure 6). In addition, DHA is the active metabolite of artemisinin, another of our cross-species confirmed leads that was shown to have neuroprotective activity in rat models of stress-induced neuronal damage and light-induced photoreceptor degeneration (Yan et al., 2017). Unfortunately, we did not observe increased photoreceptor survival in *rd10* retinas injected with PLGA encapsulated DHA. One possible explanation is that *rd1* and *rd10* models have differential responses to DHA. This has been reported for the histone-deacetylase inhibitor valproic acid, which shows opposing effects in *rd1* (neuroprotective) *rd10* (deleterious) mice (Mitton et al., 2014) and across four different frog models of RP (Vent-Schmidt et al., 2017). In addition valproic acid has produced inconsistent results in clinical trials with RP patients (Chen et al., 2019; Todd and Zelinka, 2017; Totan et al., 2017). These results emphasize the need for a more thorough understanding of IRD and RP disease mechanisms and downstream cell pathways to support the development of both personalized and pan-disease therapeutics.

Numerous IRD/RP-linked mutations have been identified (Dias et al., 2017; https://sph.uth.edu/retnet/) implicating an array of disease mechanisms (Dharmat et al., 2020). However, cell death pathways common across different IRD/RP patient subpopulations may provide pan-disease targets for neuroprotective therapies. Apoptosis has long been thought to be the mechanism by which rod photoreceptors die in IRD/RP (Chang et al., 1993; Doonan et al., 2003; Portera-Cailliau et al., 1994; Zeiss et al., 2004). However, these reports relied on terminal deoxynucleotidyl transferase dUTP nick end labeling (TUNEL), which does not distinguish apoptosis from other types of cell death (Ansari et al., 1993; Charriaut-Marlangue and Ben-Ari, 1995; Grasl-Kraupp et al., 1995; Dmitrieva and Burg, 2007; Kanoh et al., 1999; Nishiyama et al., 1996). More recent evaluations of multiple apoptosis-related markers (e.g., BAX, cytochrome c, caspase-9, cleaved caspase-3) suggest apoptosis occurs in only a minority of RP models (Arango-Gonzalez et al., 2014; Sancho-Pelluz et al., 2008). Moreover, inhibition of apoptosis does not block cell death in many mouse photoreceptor degeneration models (Hamann et al., 2009; Yoshizawa et al., 2002), suggesting other pathways may mediate rod and/or cone cell death in retinal degenerative disease.

Recently, non-apoptotic cell death mechanisms have been implicated in IRD/RP. In a comprehensive biochemical analysis of ten mammalian RP models—involving mutations in *cnga3*, *cngb1*, *pde6a*, *pde6b*, *pde6c*, *prph2*, *rho*, and rpe65—non-apoptotic cell death signatures were found to be common across all models tested (Arango-Gonzalez et al., 2014). Conversely, definitive apoptotic markers were found only for the S334ter (*rho*) rat model. Shared features included activation of poly-ADP-ribose-polymerase (PARP), histone deacetylase (HDAC), and calpain, and accumulation of cyclic guanosine monophosphate (cGMP) and poly-ADP-ribose (PAR). For PARP, chemical inhibitors and a knock out line provided further confirmation (Jiao et al., 2016; Paquet-Durand et al., 2007; Sahaboglu et al., 2017, 2016, 2010). Prior reports had suggested the NTR/Mtz system elicits caspase-3 activation and apoptotic cell death (Chen et al., 2011). Initially, we had used this as a rational for pursuing a neuroprotective screen with the *rho:YFP-NTR* line, however, the recent reports outlined above suggested apoptosis may have limited relevance to IRD/RP. Interestingly, when we tested cell death processes implicated in photoreceptor degeneration directly, an inhibitor of apoptosis (BEL) did not promote rod cell survival whereas inhibition of necroptosis (NEC) was neuroprotective in our fish RP model (Figure 4, Table 2). Serendipitously, an exploration of shared lead compound MOA helped to clarify cell death mechanism(s) mediating NTR/Mtz-induced rod cell ablation.

To explore molecular MOA of our lead compounds, we searched bioactivity data from prior HTS and ultra HTS assays (PubChem). The results suggested both shared and independent MOA (Supplementary Table 5). The most common shared target was TDP1 (Supplementary Table 5, eight of eleven lead compounds). An initial test confirmed two of three TDP1 inhibitors (Figure 4). TDP1 is a DNA repair enzyme that repairs topoisomerase I-induced DNA damage (Dexheimer et al., 2008; El-Khamisy, 2011). Interestingly, a qHTS cell-based screen of 400,000 compounds for inhibitors of human TDP1 found that all five confirmed compounds actually inhibited PARP activity not TDP1 (Murai et al., 2014). This is consistent with findings showing that TDP1 acts in conjunction with PARP1 (Das et al., 2014; Lebedeva et al., 2015), thus PARP inhibition can indirectly affect TDP1 activity. Combined with the results discussed above, we were motivated to test whether PARP inhibition was protective against NTR/Mtz-induced rod cell death. All eight PARP inhibitors tested promoted rod cell survival in our fish RP model (Figure 4 and Table 2). This result suggests our lead compounds may promote rod cell survival primarily by inhibiting a PARP-dependent cell death pathway.

PARP1 overactivation initiates a caspase-independent form of DNA-damage induced cell death pathway, termed parthanatos (Fan et al., 2017). Parthanatos has been strongly implicated in Parkinson’s disease (Kam et al., 2018) and is involved in a variety of neurodegenerative conditions as well (Fan et al., 2017). For instance, PARP has been identified as a common factor across a series of mammalian retinal disease models spanning all major forms of hereditary human blindness (Arango-Gonzalez et al., 2014). Importantly, PARP is also a key component of the cGMP-dependent cell death pathway which has been linked to photoreceptor degeneration (Iribarne and Masai, 2017; Tolone et al., 2019; Power et al., 2019). As noted above, we found that inhibiting necroptosis but not apoptosis promoted rod cell survival in fish (Figure 4 and Table 2). Necroptosis is primarily associated with secondary cone cell death in RP models (Murakami et al., 2015, 2012; Yang et al., 2017) but has also been implicated in rod cell death in *IRBP* mutant RP models (Sato et al., 2013) and/or may damage photoreceptors indirectly via necroptotic microglia signaling (Huang et al., 2018). Combined, these results suggest that necroptosis, PARP1-dependent parthanatos, and/or c-GMP-dependent cell death mediate NTR/Mtz-induced rod cell ablation. The potential relevance of these alternative cell death pathways to heritable photoreceptor degeneration may explain the relatively high rate of validation we observed in cross-species tests of lead compounds. PARP1 has also been shown to have a role in stem cell reprogramming (Chiou et al., 2013; Doege et al., 2012; Weber et al., 2013). PARP inhibition might therefore block Müller glia dedifferentiation, diverting injury-induced activity from a regenerative to neuroprotective program (Bringmann et al., 2009). That all four compounds which inhibited rod cell regeneration (MIC, CLO, CPO and DHA) were also implicated as PARP inhibitors is consistent with this possibility.

Finally, MOA analyses also revealed numerous independent lead compound targets (Supplementary Table 5). Compounds acting through independent targets/pathways have the potential to produce additive or even synergistic effects. This possibility was confirmed for 10 of 19 viable paired lead compound assays (Figure 7, Supplementary Figure. 9). This result exemplifies a key advantage of phenotypic screening: the potential to identify multiple signaling pathways providing inroads to the desired therapeutic endpoint. That combinatorial assays could be performed efficiently and rapidly also exemplifies key advantages the zebrafish system affords phenotypic drug discovery, e.g., versatility and low cost.

In summary, we identified eleven lead compounds promoting rod cell survival in an inducible zebrafish RP model using a large-scale *in vivo* phenotypic drug discovery platform. Seven lead compounds were also effective as neuroprotectants in either primary mouse retinal cell cultures or in *rd1* mouse retinal explants. MOA studies indicated lead compounds may protect rod cell from death by inhibiting PARP-dependent cell death pathways and/or necroptosis. Combinatorial assays in fish showed additive effects, suggesting lead compounds also target independent neuroprotective pathways (summarized in Figure 7). We hypothesize that compounds producing beneficial outcomes across diverse animal disease models likely target highly conserved MOA and may therefore stand an increased likelihood of successfully translating to the clinic. Further, our data suggest polypharmacological targeting of complementary neuroprotective mechanisms has the potential to maximize therapeutic benefits for IRD/RP patients.

## MATERIALS and METHODS

All animal studies described herein were performed in accordance with both the Association for Research in Vision and Ophthalmology (ARVO) statement on the “Use of Animals in Ophthalmic and Vision Research” and the National Institutes of Health (NIH) Office of Laboratory Animal Welfare (OLAW) policies regarding studies conducted in vertebrate species. Animal protocols were approved by the Animal Care and Use Committees of the Johns Hopkins University School of Medicine and Medical University of South Carolina.

### Fish maintenance and husbandry

Zebrafish were maintained using established temperature and light cycle conditions (28.5°C, 14 hr of light/10 hr of dark). A previously generated zebrafish transgenic line, *Tg(rho:YFP-Eco.NfsB)gmc500* (hereafter, *rho:YFP-NTR*) expresses a fusion protein of enhanced yellow fluorescent protein (YFP) and a bacterial nitroreductase (NTR) enzyme (encoded by the *E. coli nfsB* gene) selectively in rod photoreceptors (Walker et al., 2012). When *rho:YFP-NTR* fish are exposed to the prodrug metronidazole (Mtz), NTR reduces Mtz to a DNA damage-inducing cytotoxic derivative, resulting in the death of NTR-expressing cells (Curado et al., 2007; White and Mumm, 2013). This line was propagated in a pigmentation mutant with reduced iridophore numbers, *roy^a9^* (*roy*), to facilitate YFP reporter signal detection *in vivo*. Non-transgenic *roy* fish were used to define reporter signal cutoff values for fluorescent microplate reader assays. For each drug assay, 6,000-12,000 eggs were collected by group breeding ∼300 adult *rho:YFP-NTR* fish (White et al., 2016).

### Immunostaining

Zebrafish *Tg*(*rho:YFP-NTR)* larvae were treated with 2.5 mM Mtz or 0.1% DMSO at 5 dpf for 2 days, and collected at 7 dpf for immunostaining using previously reported methods (Unal Eroglu et al., 2018). Briefly, larvae were fixed in 4% PFA at 4°C overnight, infiltrated with 30% sucrose, and embedded in OCT. Cryosections of 10 µm thickness were made for immunofluorescence staining. Each section was blocked with 1x PBS containing 0.5% Triton X-100 and 5% goat serum for one hour followed by primary antibody staining overnight. The next day, each section was washed and stained with the secondary antibody for 2 hours. All slides were mounted and underwent confocal imaging. The primary antibodies used in this study were zpr1 (anti-arrestin3a; 1:200; from ZIRC), 1d1 (anti-Rho; 1:50; gift from Dr. James M. Fadool) and anti-4c12 (1:100; gift from Dr. James M. Fadool). The secondary antibody used was goat anti-mouse Alexa 647 (1:1000; Invitrogen).

### Optimization of Mtz concentration to establish an inducible RP model

*rho:YFP-NTR* larvae were separated into six groups of 56 larvae per group. Each group was treated with varying Mtz concentrations (10, 5, 2.5, 1.25, 0.625, 0 mM), from 5-8 days post-fertilization (dpf). YFP signals were quantified daily by fluorescence microplate reader (TECAN Infinite M1000 PRO; excitation 514 nm, bandwidth 5 nm; emission 538 nm, bandwidth 10 nm) to track changes in rod photoreceptor cell numbers relative to Mtz concentration (Walker et al., 2012; White et al., 2016).

### Sample size estimation

To calculate sample size (n) and evaluate assay quality, two 96-well plates of larvae were treated with 2.5 mM Mtz/0.1% DMSO (ablated “+Mtz” control) or 0.1% DMSO only (non-ablated “-Mtz” control) from 5-7 dpf. YFP signals were then quantified at 7 dpf. Power calculations were used to determine sample sizes across a range of error rates and effect sizes (White et al., 2016). This analysis suggested a sample size of nine per condition tested would sufficiently minimize false-discoveries (i.e., type I and type II error rates of 0.05 and 0.05, respectively) and account for drug effects reaching 50% of signal window of the positive (non-ablated) control. However, to account for plating errors and other confounding variables, we chose to increase the sample size to 16.

### Primary drug screening using ARQiv platform

A schematic of the primary screening process is presented in Fig. 1C. For the primary screen, 2,934 compounds from John Hopkins Drug Library (JHDL) were screened across six concentrations (4 μM-125 nM using a 2-fold dilution series). The JHDL consists of ∼2,200 drugs approved for use in humans (e.g., FDA approved) with the remainder approved for clinical trials (Chong et al., 2006). The ARQiv screening process has been detailed previously (White et al., 2016) and was adapted here for large-scale quantification of YFP-expressing rod photoreceptors (Walker et al., 2012; White et al., 2016). *rho:YFP-NTR* embryos were collected and raised in zebrafish E3 embryo media (5 mM NaCl; 0.17 mM KCl; 0.33 mM CaCl; 0.33 mM MgSO_4_). At 16 hours post fertilization (hpf), N-phenylthiourea (PTU) was added to E3 media (E3/PTU) at a final concentration of 0.2 mM to promote ocular transparency by inhibiting melanosome maturation in the retinal pigment epithelium. At 4 dpf, visual screens were performed to remove larvae with abnormal morphology or low YFP expression levels. Stock drug and DMSO (negative control) solutions were automatically dispensed and diluted across a 96-well plate containing E3/PTU using a robotic liquid handling system (Hudson Robotics). At 5 dpf, a COPAS-XL (Complex Object Parametric Analyzer and Sorter, Union Biometrica) was used to dispense single larvae into individual wells containing either drug or DMSO; the final DMSO concentration was 0.1% across all conditions. After a four hr pre-exposure to test drugs or DMSO alone, larvae were treated with 2.5 mM Mtz to induce rod photoreceptor death. Larvae were maintained under these conditions for two days until 7 dpf and then anesthetized with clove oil (50 ppm final concentration); YFP signals were measured as described above. Larvae exposed to 2.5 mM Mtz/0.1% DMSO without any tested drug served as controls (“+Mtz”) for maximal rod cell ablation. Larvae treated solely with 0.1% DMSO served as non-ablated controls (“-Mtz”) to calculate maximal YFP signal levels. Non-transgenic larvae were used to establish a signal cutoff value, as previously described (White et al., 2016). A customized R program was applied for real-time data analysis of compound performance relative to controls, including dose-response curves and strictly standardized mean difference (SSMD) scores. Compound concentrations producing a SSMD score ≥1 were considered potential ‘hits’ and evaluated visually using fluorescence microscopy to eliminate non-specific fluorescence; e.g., dead larvae or autofluorescent compounds.

### Secondary validation tests of potential hits

After the initial screen, 42 top-performing ‘hit’ compounds were selected for confirmatory and orthogonal assays. Hit drugs were obtained from new sources and tested across a wider range of concentrations (five-fold dilution series, eight total concentrations, n=30 fish/condition). Based on the toxicity profile of each drug, the starting concentration was either 100, 10 or 1 mM. For validation assays, YFP signals were measured by fluorescence microplate reader with three biological replicates conducted and SSMD scores calculated. All experimental results were normalized and pooled to calculate effect sizes, confidence intervals, and *p*-values using Student’s t-test followed by a Bonferroni corrected *p*-value criterion of 0.005 (α/n).

### Confocal imaging

For *in vivo* confocal imaging assays, 5 dpf *rho:YFP-NTR* larvae were treated with drug plus 2.5 mM Mtz/0.1% DMSO (ablated +Mtz controls), or 0.1% DMSO (non-ablated - Mtz controls) for 48 hrs and processed for intravital imaging at 7 dpf. All compounds were tested at the maximal effective concentration in 0.1% DMSO. Larvae from each group were anesthetized in 0.016% tricaine and embedded on their sides in 1% low melt agarose gel. An Olympus Fluoview FV1000 confocal microscope with a 20x water immersion objective (0.95 NA) was used to collect 30-40 images of YFP-expressing rod cells across the whole retina at 4 μm intervals. These image slices were stacked into a single maximal intensity projection image using Image J. A region in the dorsal-nasal quadrant was imaged using a 60× water immersion objective (1.10 NA) at 4.18 μm intervals to provide greater detail of rod photoreceptor morphology. Three contiguous image slices were stacked into a single maximal intensity projection image using Image J. As a means of assessing rod photoreceptor survival, intravital confocal retinal images were collected of YFP-expressing rod cells per condition and processed for volumetric quantification of YFP signal volume using automated image analysis software (Imaris 3.9.1, see White et al., 2017). Briefly, YFP-expressing rod cells volumes were automatically rendered using the same parameters across all treatment groups with the sum of all volumes used to estimate rod photoreceptor numbers per each retina. Sample sizes for rod cell quantification assays ranged from 6 to 13 per condition.

### Nitroreductase inhibitory assay

The eleven lead compounds were tested to eliminate false-positives that directly inhibited *Escherichia coli* NTR enzymatic activity (from the *nfsB_Ec* gene) using an established anticancer prodrug (CB1954) reduction assay (Prosser et al., 2010). The highest soluble concentration of the test drug (up to 300 M maximum) or 0.1% DMSO (rate control) was added to a master mix of 250 μM CB1954/1 μM NTR/250 μM NADPH/10 mM Tris (pH 7.0) to determine any inhibitory effect of the compounds on NTR-mediated reduction of CB1954. Reactions were conducted in 100 μl volume in a 96-well plate format. CB1954 reduction kinetics were assayed at 420 nm unless the test compound exhibited confounding absorbance at 420 nm, in which case NADPH depletion was monitored at 340 nm. The reaction rates for CB1954 reduction in the presence of tested compounds were compared to the DMSO control. Compounds with NTR activity <75% of controls were deemed potentially inhibitory and IC_50_ values (the concentration required to restrict NTR activity to 50% of the DMSO control) determined.

Compounds were also evaluated for the ability to inhibit NTR-mediated reduction of Mtz. For this assay, the highest soluble concentration of the test drug (up to 300 μM maximum) or DMSO (rate control) was added to a master mix of 500 μM Mtz/17 μM NTR/100 μM NADPH/0.55 μM glucose dehydrogenase/5 mM glucose/50 mM sodium phosphate buffer (pH 7.0; the glucose dehydrogenase used for cofactor regeneration, maintaining a steady-state of NADPH to allow Mtz reduction to be monitored directly at 340 nm without interference from NADPH oxidation). Reactions were conducted in 100 μl volume and assayed at 340 nm using 1 mm quartz cuvettes (Rich et al., 2018). IC_50_ values for potentially inhibitory compounds were measured as with CB1954.

### Rod photoreceptor neogenesis assay

To evaluate whether compounds promoted rod photoreceptor development, *rho:YFP-NTR* larvae were handled as described for the primary screen with the following exceptions: at 5 dpf, larvae were exposed solely to test compounds; larvae were not treated with Mtz. YFP reporter signals were quantified by fluorescence microplate reader at 7 dpf and compared to non-ablated “-Mtz” controls treated only with 0.1% DMSO. Sample sizes ranged from 58 to 88 for lead compounds.

### Rod photoreceptor regeneration assay

To determine whether compounds stimulated retinal regeneration, *rho:YFP-NTR* larvae were handled as described for the primary screen with the following exceptions: at 5 dpf, larvae were incubated with either 10 mM Mtz or 0.1% DMSO for 24 hrs. At 6 dpf, larvae were then placed in new 0.1% DMSO/E3/PTU media containing test compounds (or DMSO alone) for three days. YFP signal intensity was measured at 9 dpf. Sample sizes ranged from 31 to 83 for lead compounds.

### Inhibitor assays for assessing mechanism of cell death

To evaluate potential shared MOA, and other cell death pathways implicated in photoreceptor degeneration, nine PARP inhibitors, one necroptosis inhibitor and one apoptosis inhibitor were selected for testing using the primary neuroprotectant screen protocol. Based on the toxicity profile of each drug, upper concentrations were adjusted to 64, 32, or 8 µM. A two-fold dilution series across a total of ten concentrations was then tested, at a sample size of 16 fish per condition. YFP signal levels were measured by fluorescence microplate reader (i.e., ARQiv assay). Results across conditions were normalized to non-ablated controls and pooled across experimental repeats to calculate effect sizes, confidence intervals, and *p*-values using Student’s t-test followed by a Bonferroni corrected *p*-value criterion of 0.005 (α/n). Note: assessments of TDP1 inhibitors followed the five-fold dilution series protocol used for secondary confirmation assays described above.

### In vitro assay for the protection of mouse retinal cells from exogenous stressors

Mice were housed with 12 hr light/12 hr dark cycles, at 22°C, 30–70% relative humidity and food/water *ad libitum*. Primary mouse retinal cells were isolated and prepared for culture as previously described (Fuller et al., 2014). Briefly, murine retinas were isolated at postnatal day four (P4). Retinal tissue was dissociated into a single cell suspension by incubating whole tissue in activated papain in Hibernate-E without Calcium (BrainBits) for 15 min at 37°C. Cells were resuspended in culture media (Neurobasal, 2% B-27, 0.5 mM L-Glutamine and 1x final Penicillin/streptomycin; all Life Technologies) and seeded onto poly-D-lysine coated 384 well tissue culture plates. Tunicamycin and thapsigargin were used as stressor compounds (Sigma). Stressors as well as test compounds were added at the time of seeding. After 48 hours, cell viability was assessed using CellTiterGlo, a single step chemiluminescent reagent that measures ATP as a proxy for relative cell survival. To assess protection specifically of photoreceptors, cells harvested from transgenic mice that contain the human QRX locus with an IRES-GFP cassette, in which GFP expression is restricted to the photoreceptor cell population, were cultured as described above. For these assays to assess photoreceptor survival, cells are stained with Hoescht and ethidium homodimer; nine field images were acquired via an automated imager (Cellomics Vti; ThermoFisher) using a 20x objective and images analyzed. Photoreceptor number and percent of retinal cell population per well were determined by quantifying the number of live Hoechst-stained, GFP-expressing cells using a custom algorithm (Neuronal Profiling software package; ThermoFisher).

### Retinal explant assay using Pde6b^rd1^ (rd1) mouse

Mice were generated from *retinal degeneration 1* (*Pde6brd1*, hereafter *rd1*) breeding pairs (Dr. Debora Farber; UCLA) and housed under a 12:12 light:dark cycle with access to food and water *ad libitum*. All chemicals used for organ cultures were tissue culture grade and purchased from Invitrogen unless otherwise noted. Cultures of retina with attached retinal pigment epithelium (RPE) were grown according to published protocols (Ogilvie et al., 1999; Pinzón-Duarte et al., 2000; Rohrer and Ogilvie, 2003) with modifications (Bandyopadhyay and Rohrer, 2010). P10 pups were decapitated and heads were rinsed in 70% ethanol; whole eyes were dissected and placed in ice-cold Hanks balanced salt solution plus glucose (6.5 g/L). Eyes were first incubated in 1 mL of high glucose Hanks balanced salt solution/0.5 mg/ml proteinase K (37°C, 7 min) and then in Neurobasal medium (Life Technologies) plus 10% fetal calf serum to stop enzymatic activity. The retina with attached RPE was dissected free from the choroid and sclera after removing the anterior chamber, lens and vitreous. Relaxing cuts were made to flatten the tissue prior to transfer to the upper compartment of a Costar Transwell chamber (RPE layer faced-down) using a drop of Neurobasal medium. Neurobasal media with B-27 supplement (Life Technologies) was placed in the lower compartment. For each of the cohorts, 3-4 individual P10 retina/RPE explants were placed in culture. The cultures were kept in an incubator (5% CO_2_, balanced air, 100% humidity, 37°C) and the lower compartment media changed every two days (neither antimitotics nor antibiotics were required). Test compounds were added to the culture media and refreshed every two days for 11 days. At completion, explants were fixed in 4% PFA, sectioned 14 µm thick, and stained with HE (Bandyopadhyay and Rohrer, 2010). For each culture, ten rows of photoreceptors along the length of each culture were counted; ten values were averaged to give a value for each retina, while the average of all retinas provided the mean ± SEM of each culture condition. ANOVA test was performed across all groups followed by Bonferroni/Dunn t-test with significance level cutoff of p≤0.05.

### DHA long-release formulation

PLGA-DHA microparticles were prepared using a single emulsion solvent evaporation method. Briefly, 200 mg PLGA (2A, 50:50 LA:GA) (Evonik Corporation, Piscataway, NJ) was dissolved in 1 mL of dichloromethane (DCM, Sigma-Aldrich), and mixing with 40 mg DHA (TCI, Tokyo, Japan) dissolved in 0.125 ml dimethyl sulfoxide (DMSO, Sigma-Aldrich). The mixture was homogenized (L4RT, Silverson Machines) at 7000 RPM for 1 minute. The homogenized mixture was then poured into a solution containing 1% polyvinyl alcohol (25 kDa, 88% hydrolys, Polysciences, Warrington, PA) in water under continuous stirring. Particles were hardened by allowing solvent to evaporate while stirring at room temperature for 2 h. Particles were collected via centrifugation (International Equipment Co) at 1,000 x g for 5 min, and washed three times with HyPure cell culture grade water (endotoxin-free, HyClone™, Logan, UT) and re-collected by centrifugation three times. The washed particles were then lyophilized and stored frozen until use. Microparticles were resuspended at the desired concentration prior to injection in a sodium hyaluronate solution (Healon®) diluted 5-fold with endotoxin-free HyPure water at the desired concentration prior to injection.

### Characterization of PLGA-DHA microparticles

Particle size distribution was determined using a Coulter Multisizer 4 (Beckman Coulter, Inc., Miami, FL). Particles were resuspended in double distilled water and added dropwise to 100 ml of ISOTON II solution until the coincidence of the particles was between 8% and 10%. At least 100,000 particles were sized for each batch of particles to determine the mean particle size and size distribution. To determine the drug loading, microparticles were dissolved in DMSO and the total drug content was calculated by measuring the UV absorbance at 289 nm after react with NaOH in triplicate (C.-S. Lai et al., 2007). The average microparticle size was 14.2 ± 1.9 µm and the DHA loading was 20.3% (w/w).

### DHA release

Release kinetics were obtained by resuspending microparticles in 1 ml phosphate buffered saline (PBS containing 0.1% DMSO, pH 7.4) and incubating at 37°C on a platform shaker (140 RPM). Supernatant was collected at predetermined intervals by centrifugation at 2,000 x g for 5 min. Drug-containing supernatant was collected and particles were resuspended in 1 ml of fresh 1xPBS containing 0.1% DMSO. DHA concentration in the collected supernatant was assayed via absorbance at 289 nm in triplicate for each sample (n=3) (Supplementary Figure 7).

### Retina injection assay using Pde6b^rd10^ (rd10) mouse

B6.CXB1-*Pde6b^rd10^*/J animals (Jackson Laboratories stock #004297) were maintained as homozygotes. On P14, animals were anesthetized with ketamine/xylazine (115 mg/kg and 5 mg/kg, respectively), and placed on a heating pad to maintain body temperature throughout injection and recovery. Glass needles were pulled and a bore size of approximately 75 µm was made to allow for the movement of the microspheres into and out of the needle. A PLI-100 picospritzer (Harvard Apparatus) was used for intravitreal injections with settings of 350 ms injection time at 30 psi, which yielded the injection of approximately 500 nL. One eye was randomly selected for injection with vehicle and the other was used for microsphere injections. Following injection, GenTeal gel (Novartis) was applied to the eyes to prevent corneal drying. Animals were monitored until they were awake and moving normally.

Eighteen days post-injection, animals were anesthetized with isoflurane and decapitated; the eyes were removed and placed in 4% PFA for 30 minutes. After removing the cornea and lens, the eye cups were placed back into fixative for 2 hrs on ice. After thorough washing with 1× PBS, eyecups were soaked in 15% sucrose (w/v) in 1× PBS for three hrs, then incubated in 30% sucrose (w/v)/1× PBS overnight. Samples were embedded in OCT compound (TissueTek) and frozen on dry ice. Cryosections of 12-14 µm thickness were cut on a Leica CM3050S cryostat and were mounted on Superfrost Plus slides (Fisher Scientific). Slides were rehydrated with 1× PBS and then incubated in blocking buffer (10% Normal donkey serum/ 0.3% Triton X-100/1× PBS) for 2 hrs, and were then incubated for 30 hrs at 4°C with primary antibodies diluted in blocking buffer. Sections were washed in 1× PBS and incubated for two hrs at room temperature with secondary antibodies diluted in blocking buffer. Following additional washes, sections were incubated with DAPI and mounted with FluoroGel (Electron Microscopy Sciences). The following antibodies were used for immunohistochemistry: Mouse mAb, anti-S-antigen S128 (Abcam 190315; 1:1000); Rabbit anti-Cone Arrestin (Millipore AB15282; 1:1000). Fluorophore-conjugated secondary antibodies were obtained from Jackson Immunoresearch, and used at a dilution of 1:1000.

Sections were examined using a Zeiss Axioplan 2 epifluorescence microscope fitted with an Axioscope camera. Following acquisition with a 40× air objective, images were analyzed in ImageJ using the measure tool to examine Outer Nuclear Layer (ONL) thickness. 4-5 measurements were taken at 75 µm intervals for 200-500 µm both superior and inferior from the optic nerve head. The measured values from multiple sections were averaged for each eye. Data were analyzed and statistical tests were performed in GraphPad Prism 7. An Olympus FV1000 confocal microscope was also used to image sections stained with multiple dyes.

### Combinatorial assay

To test for additive effects, seven lead compounds were tested at their optimal concentrations either individually or in pairs using the primary screen protocol. All experimental results were normalized to controls and pooled per compound tested to calculate effect sizes, confidence intervals, and *p*-values using Student’s t-test.Bonferroni correction for multiple comparisons resulted in an adjusted *p*-value criterion of 0.002 (α/n = 0.05/21 pairs tested).

### Data analysis and statistics

For the primary screening, under the R environment, a customized ARQiv data analysis package was used to calculate sample size (n), quality control strictly standardized mean difference (SSMD) and SSMD scores as previously described (White et al., 2016). The following results of each drug were derived: 1) a plot of signal to background ratio at all tested concentrations, 2) a plot and a table of SSMD scores, and 3) a signal intensity heat map of each drug plate (96-well plate view).

To combine and analyze the data from different experiments, data normalization was conducted by (S_i_ - *X̄_neg_*)/(*X̄_Pos_*-*X̄_neg_*). S_i_ is the signal of i^th^ reading, *X̄_Pos_* is mean of positive controls, and *X̄_pos_* is mean of negative controls. To compare between the experimental condition and controls, Student’s *t* test was performed followed by Bonferroni correction for multiple comparisons (α/n). Effect sizes, 95% confidence intervals and *p*-values were calculated.

## Acknowledgements

This project was supported by a Wynn-Gund TRAP award from the Foundation Fighting Blindness (JSM). Additional funding was provided by the Wilmer core grant for vision research, microscopy and imaging core module (NIH P30-EY001765), the National Institutes of Health (NIH) R01EY019320 (BR), the Department of Veterans Affairs RX000444 and BX003050 (BR), and the South Carolina SmartState Endowment (BR). Assembly of the JHDL was supported by the Flight Attendant Medical Research Institute and ITCR (JOL). We thank Dr. James M. Fadool for kindly providing 1d1 and 4c12 monoclonal antibodies.

## Competing interests

JSM holds patents for the NTR inducible cell ablation system (US #7,514,595) and uses thereof (US #8,071,838 and US#8431768). JSM and LZ have filed a provisional patent for the discoveries described herein. All other authors have no commercial relationships.

## Supplement Data

**Supplementary Figure 1.**
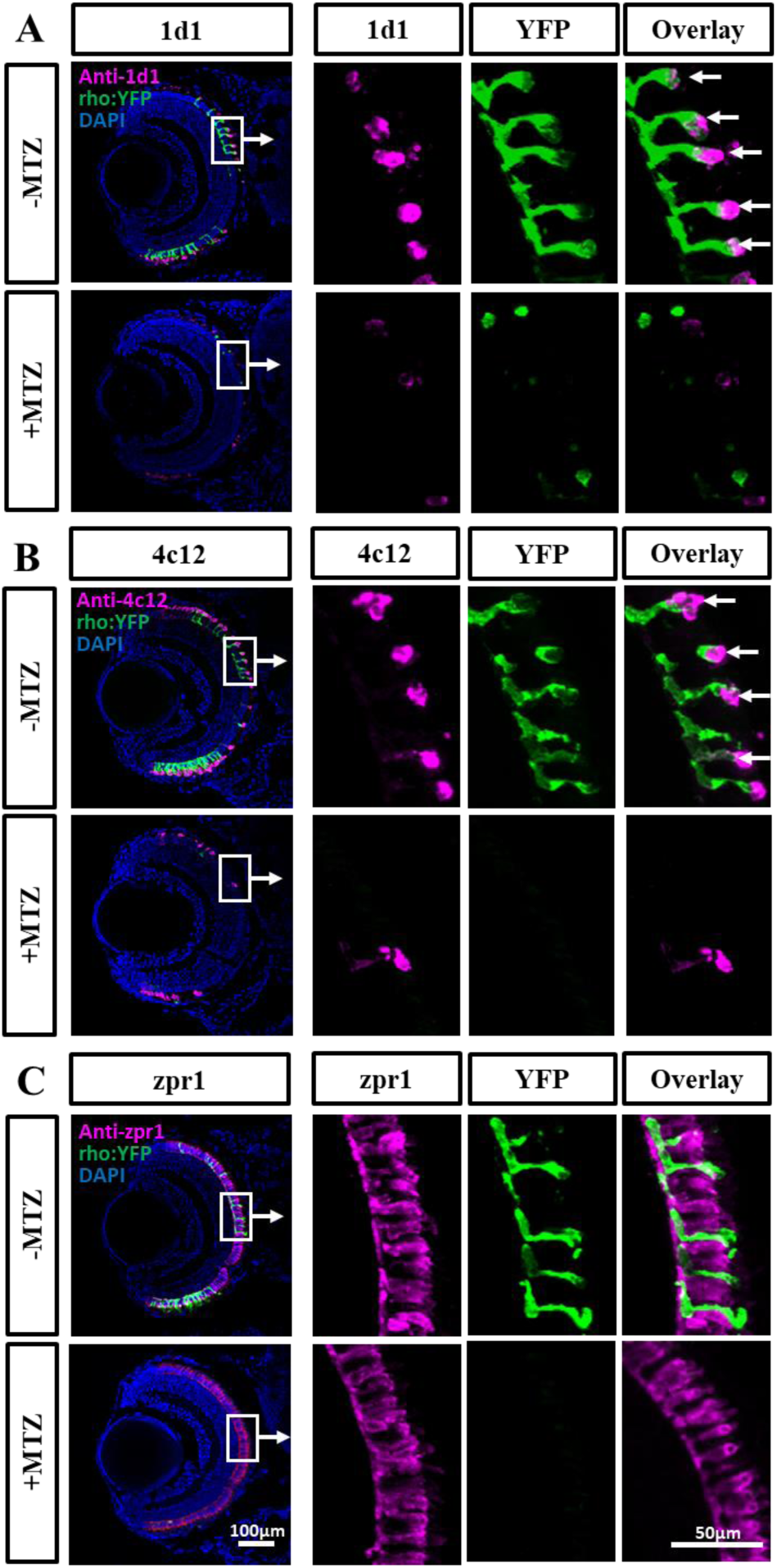
Immunolabeling of rod and cone photoreceptors. Immunohistological assays for rod and cone photoreceptor markers were performed on Mtz-ablated and control *rho:YFP-NTR* larval retinas at 7 dpf. (A) In non-ablated controls, rod photoreceptor antibody 1d1 (anti-rhodopsin) labeled rod outer segments and was well correlated with YFP expression (arrows). In Mtz-ablated retinas, both 1d1 and YFP labeling was diminished. (B) Another rod photoreceptor antibody, 4c12, showed a similar result. Both rod cell antibodies showed some residual labeling in peripheral regions in +Mtz controls. (C) Cone photoreceptors labeled with the zpr1antibody showed no overlap with YFP expression. Both Mtz-ablated and non-ablated retinas showed similar zpr1 labeling suggesting cone cells were not affected by rod cell loss. Scale bars: 50 and 100 µm.

**Supplementary Figure 2.**
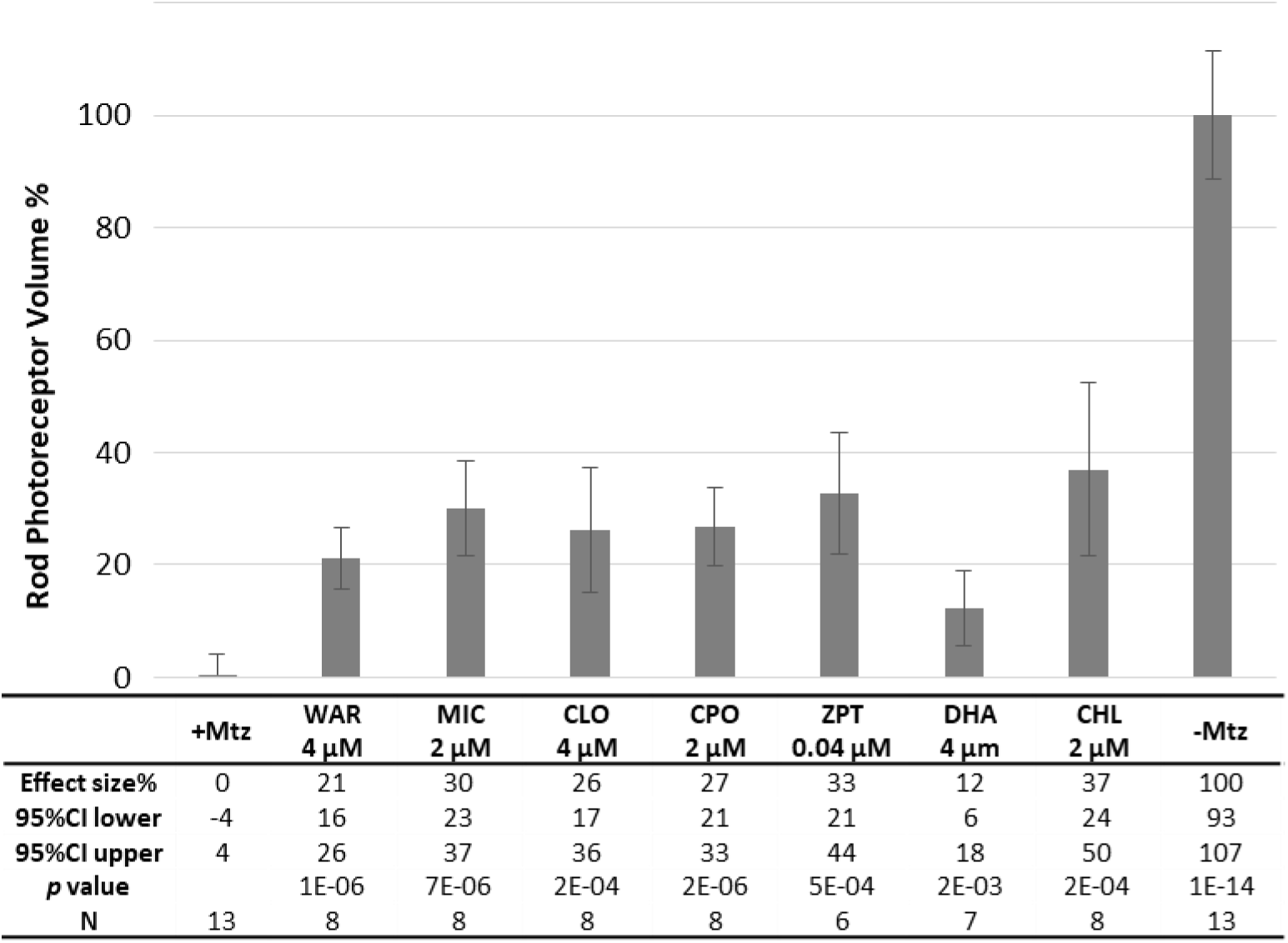
High-content imaging quantification of rod photoreceptor survival. Larvae were treated with Mtz alone (+MTZ, ablated control), Mtz plus one of seven lead compounds, or 0.1% DMSO alone (-MTZ, non-ablated control), as per the primary screen. At 7 dpf, representative retinas from drug treated and control groups were imaged using intravital confocal microscopy using identical acquisition settings and Z-scans that spanned the entire retina. Using Imaris software, YFP-expressing cells were 3D-rendered and volumetrically quantified to assess rod photoreceptor survival across all groups. The effect size, 95% confidence intervals, *p* value and sample size (N) are listed below the graph for each condition. A student’s t test followed by Bonferroni correction for multiple comparisons (α=0.006 adjusted significance level) was performed. All drug treated retinas showed a larger YFP volume, suggesting increased rod cell survival compared to the ablated group (+Mtz).

**Supplementary Figure 3.**
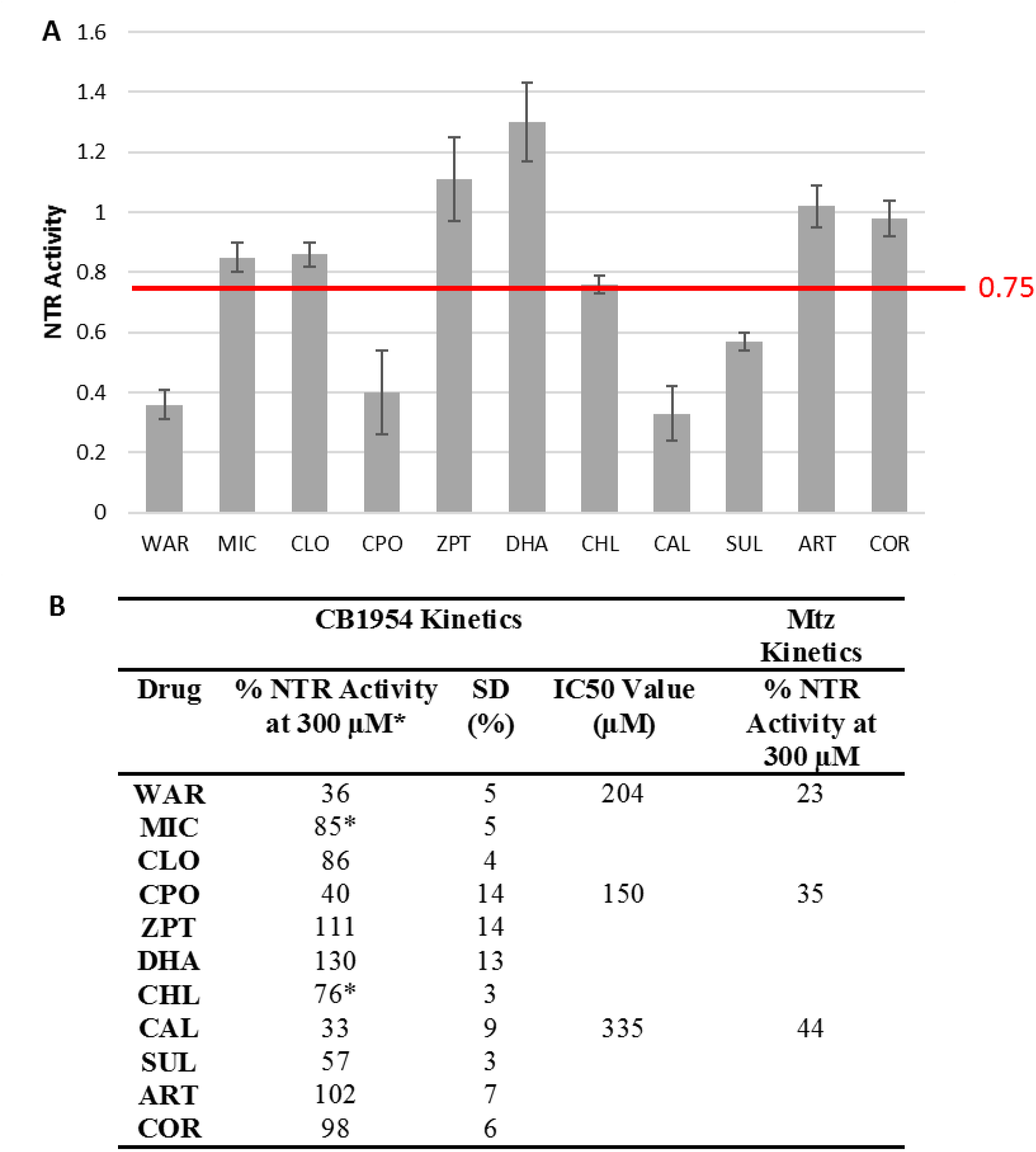
Orthogonal assay for NTR inhibition. The possibility of direct inhibition of NTR enzymatic activity by lead compounds was tested using a CB1954 kinetics assay with all leads initially tested at 300 µM with the exception of MIC and CHL which were tested at 50 µM as both precipitated at higher concentrations (indicated by *). (**A**) NTR inhibition was plotted as the ratio of NTR activity detected in drug treated samples to no-drug treated controls. Compounds producing a ratio of ≤0.75: WAR, CPO, CAL and SUL, were considered potential inhibitors. (**B**) Table showing IC_50_ of potential inhibitors for the CB1954 kinetics assay, as well as Mtz inhibition detected in a kinetics assay at 300 µM.

**Supplementary Figure 4.**
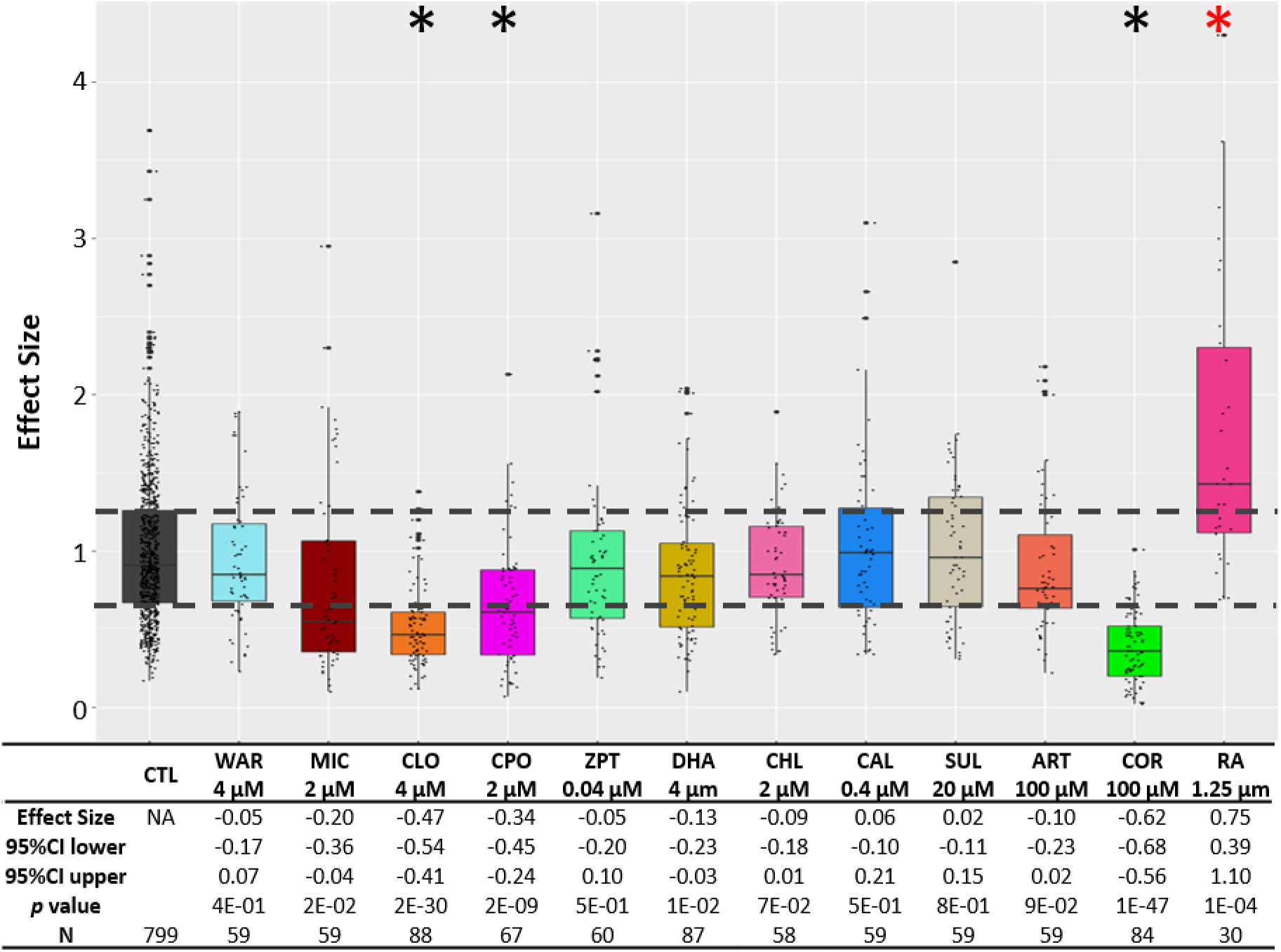
Orthogonal assay for developmental effects. Larvae were treated with individual lead compounds from 5 to 7 dpf to test for effects on rod photoreceptor development; retinoic acid (RA) served as the positive control. YFP signals were quantified by plate reader assay and normalized to DMSO-treated controls (CTL). Effect sizes, 95% confidence intervals, *p* values, and sample sizes (N) for each group are shown below. *P* values were calculated by comparing drug-treated conditions to DMSO controls using Student’s t test followed by Bonferroni correction for multiple comparisons (α=0.004 adjusted significance level). No statistical differences in larval survival were observed for lead compounds relative to their respective +Mtz controls, except for CIC (74%; Fisher’s exact test, p<0.05).

**Supplementary Figure 5.**
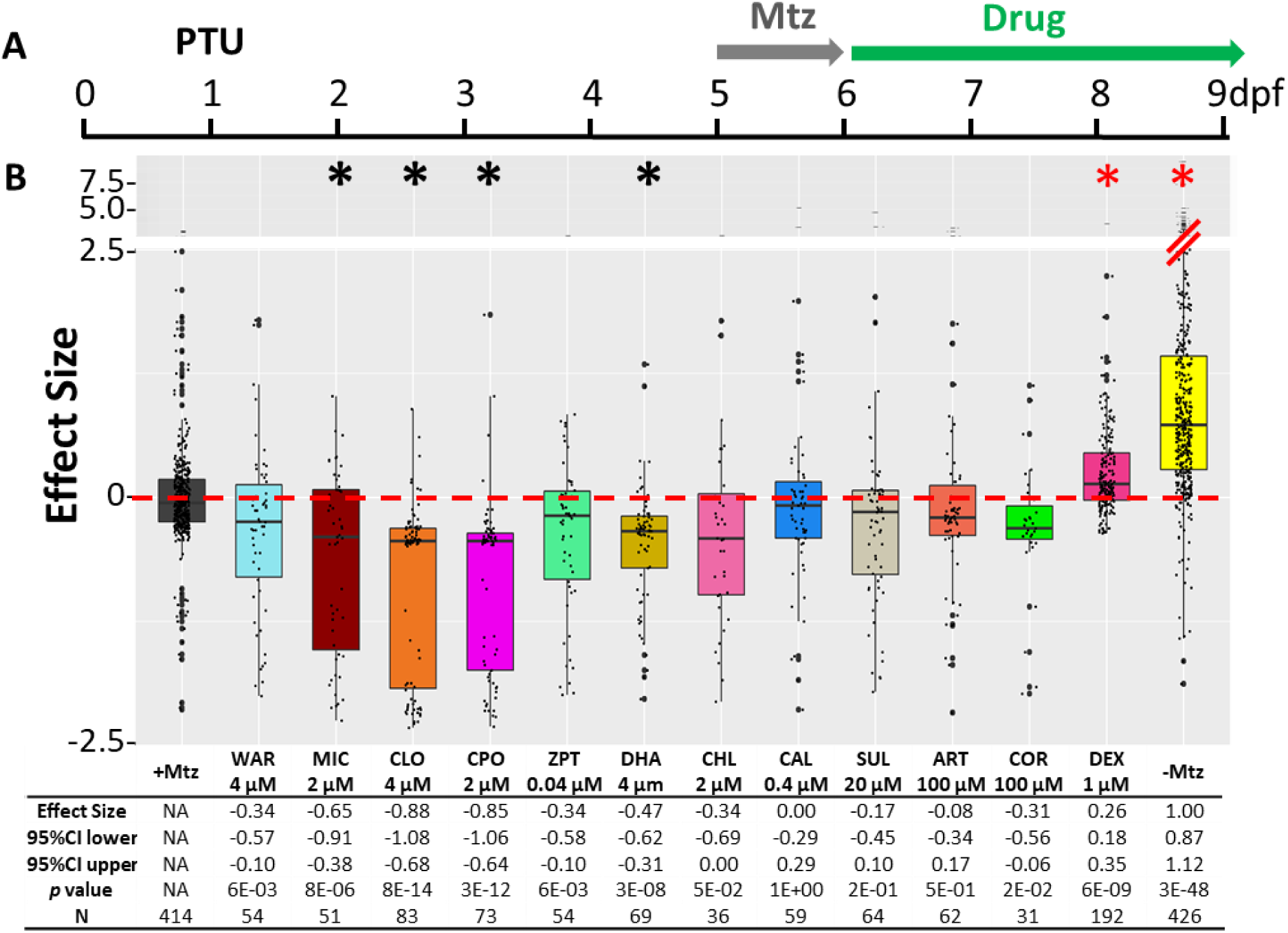
Orthogonal assay for regenerative effects. To test lead compounds for effects on rod cell regeneration kinetics, larvae were exposed to 10 mM Mtz from 5 to 6 dpf, then treated with individual lead compounds from 6 to 9 dpf; dexamethasone served as the positive control (DEX). YFP signals were quantified by plate reader assay and normalized to the signal window established by ablated (+MTZ, 0%) and non-ablated (-MTZ, 100%) controls. The effect size, 95% of confidence intervals, *p* value, and sample size (N) for each condition is shown below. *P* values were calculated by comparing drug-treated conditions to +MTZ controls using Student’s t test followed by Bonferroni correction for multiple comparisons α=0.004 adjusted significance level). Red * denotes the compounds with significant improvement effect, while black * denotes the compounds with significant reduced effect compared to DMSO-treated control. No statistical differences in larval survival were observed for lead compounds relative to their respective +Mtz controls, except for CIC (76%), DHA (72%), CHL (56%) and COR (48%; Fisher’s exact test, p<0.05).

**Supplementary Figure 6.**
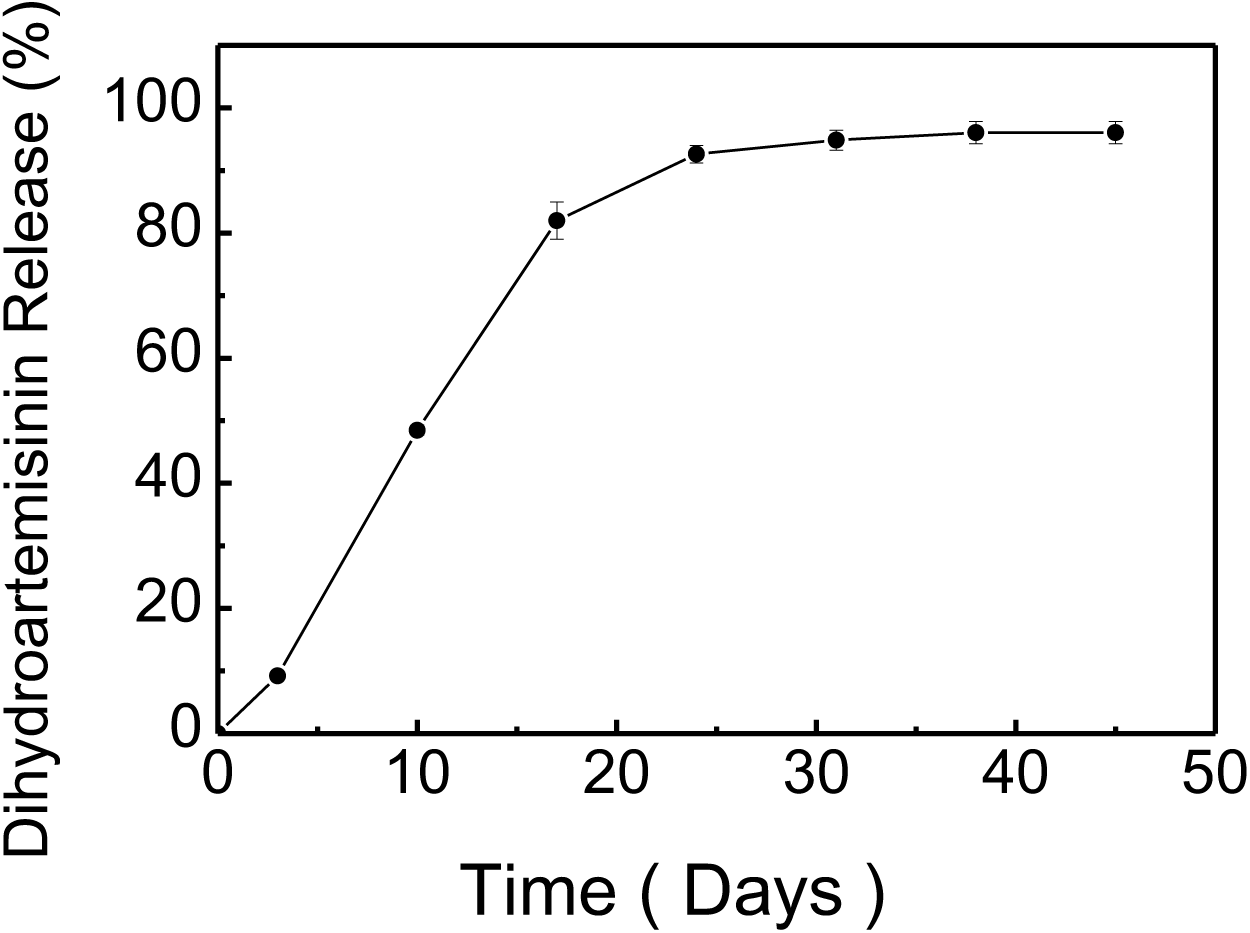
*In vitro* release kinetics of DHA from PLGA(2A) microparticles in phosphate buffered saline containing 0.1% DMSO (pH 7.4) at 37°C.

**Supplementary Figure 7.**
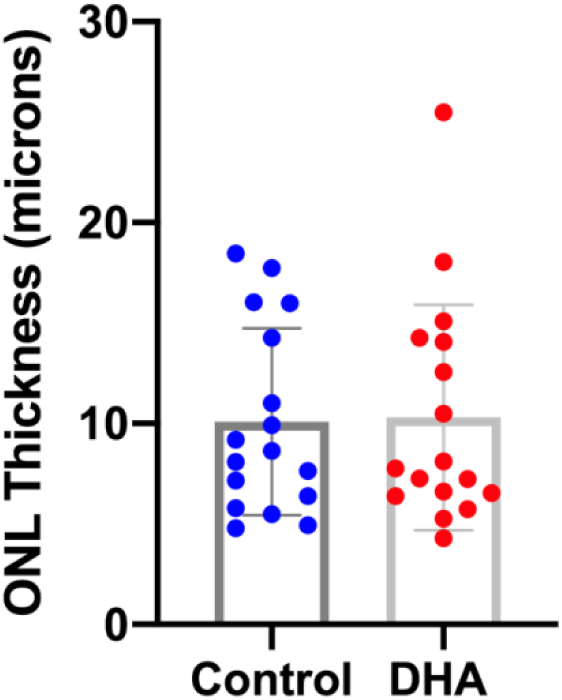
Long-release DHA evaluation in *rd10* mouse of RP. Long-release formulation of PLGA-DHA was injected into the vitreous of one eye of *rd10* mice at P14. The contralateral eye was used as a vehicle injection control. Eighteen days after injection, both eyes of each rd10 mouse were processed for immunohistochemistry. The ONL thickness was measured. No increase in ONL thickness was observed in the DHA treated group. Sample size was 17 mice.

**Supplementary Table 1.** Statistical results of Mtz titration assay using *rho:YFP-NTR* fish at 7 dpf (Figure 1A). The effect size, 95% confidence intervals, p value, and sample size (N) for each condition is listed. *P* values were calculated by comparing each condition to non-ablated controls (0mM) using Student’s t test followed by Bonferroni correction for multiple comparisons (α=0.01 adjusted significance level).

| 7dpf | 0mM | 0.625mM | 1.25mM | 2.5mM | 5mM | 10mM |
| --- | --- | --- | --- | --- | --- | --- |
| **Effect size** | NA | -0.55 | -0.79 | -0.87 | -0.87 | -0.83 |
| **95%CI lower** | NA | -0.72 | -0.95 | -1.02 | -1.02 | -0.98 |
| **95%CI upper** | NA | -0.38 | -0.63 | -0.71 | -0.72 | -0.67 |
| **p value** | NA | 9E-09 | 5E-15 | 1E-16 | 1E-16 | 1E-15 |
| **N** | 56 | 55 | 56 | 56 | 56 | 55 |

**Supplementary Table 2.**
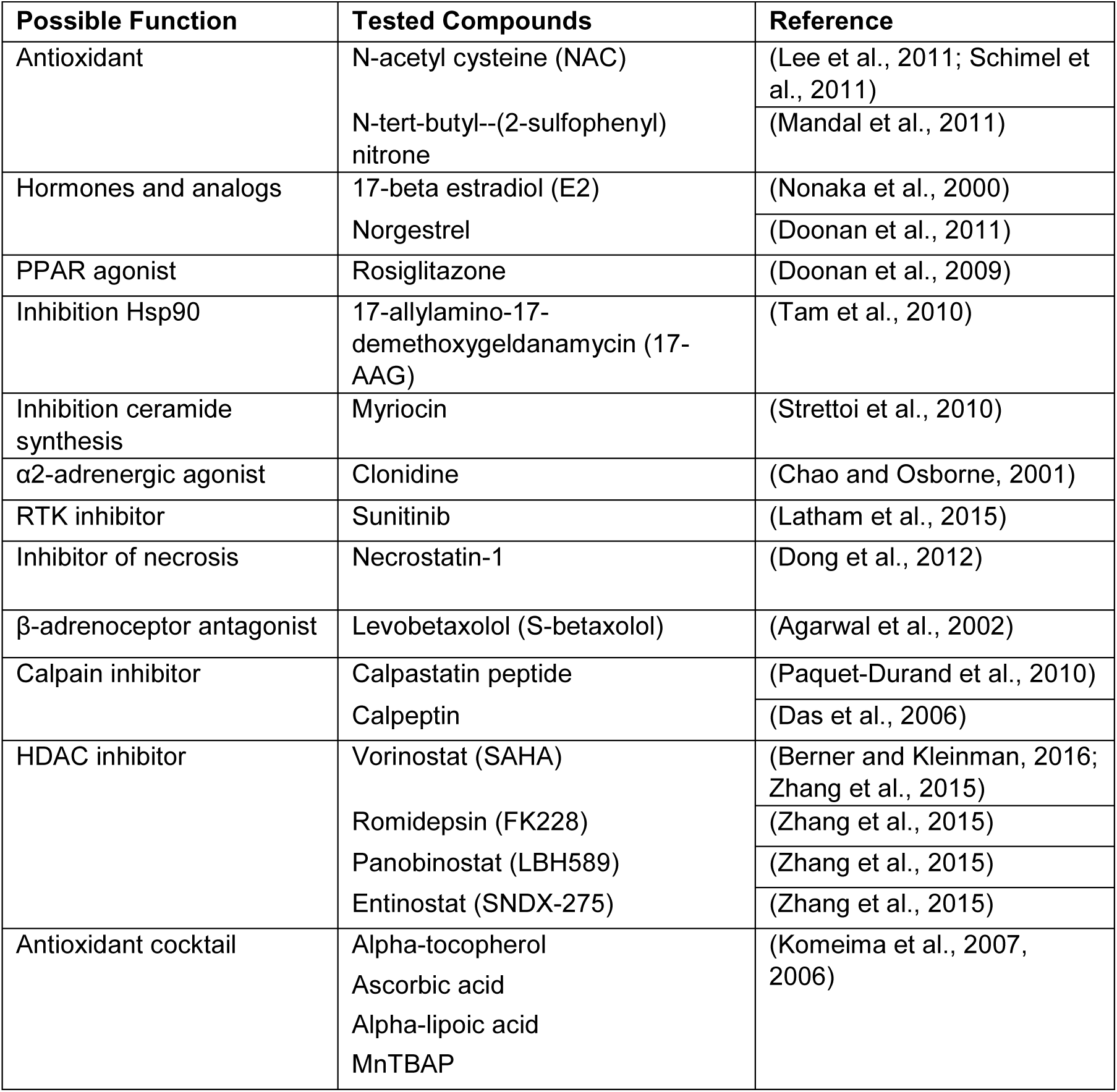
List of 17 compounds previously reported as neuroprotectants in RP models that were tested using our primary screening protocol.

**Supplementary Table 3.**
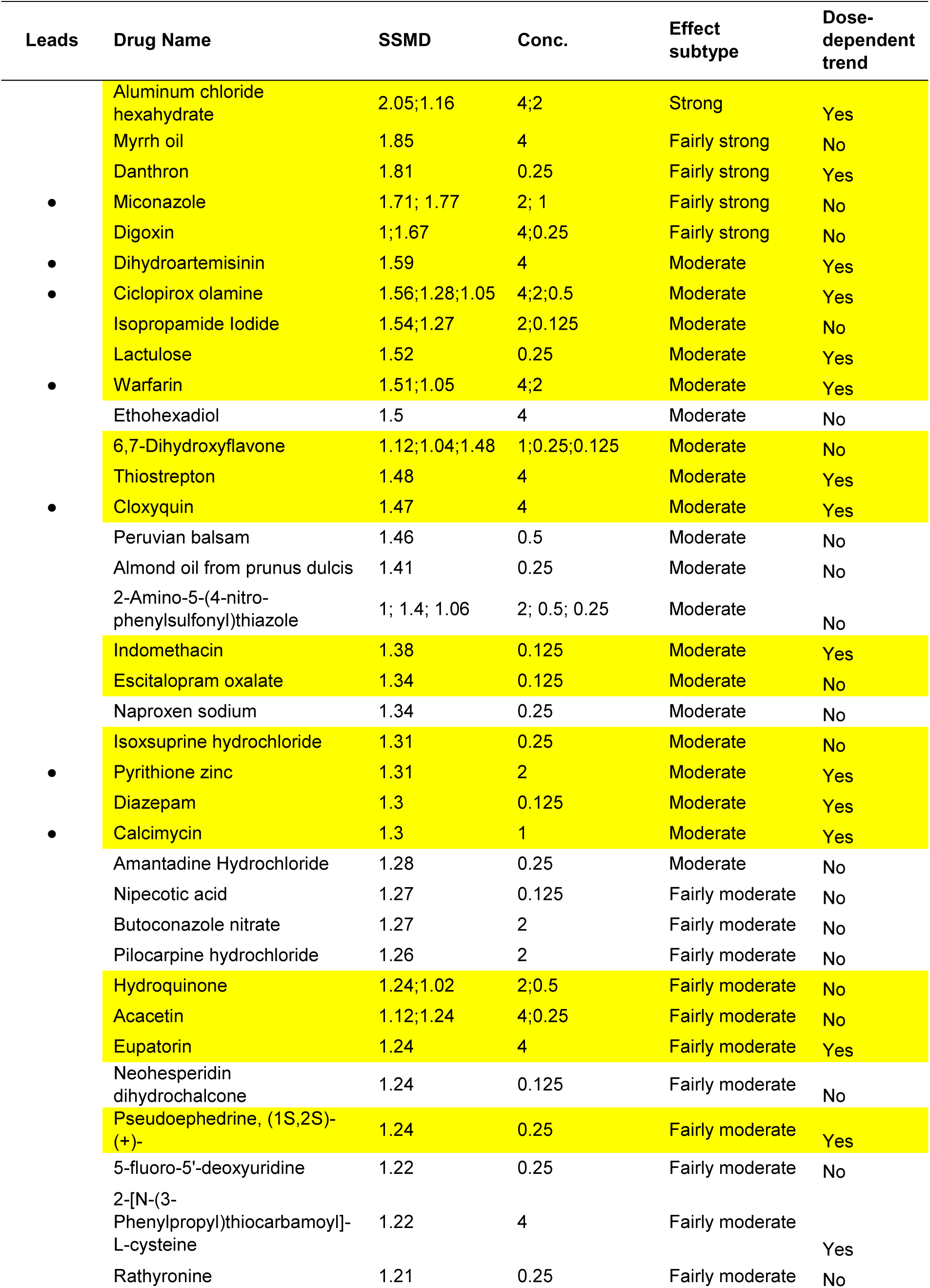
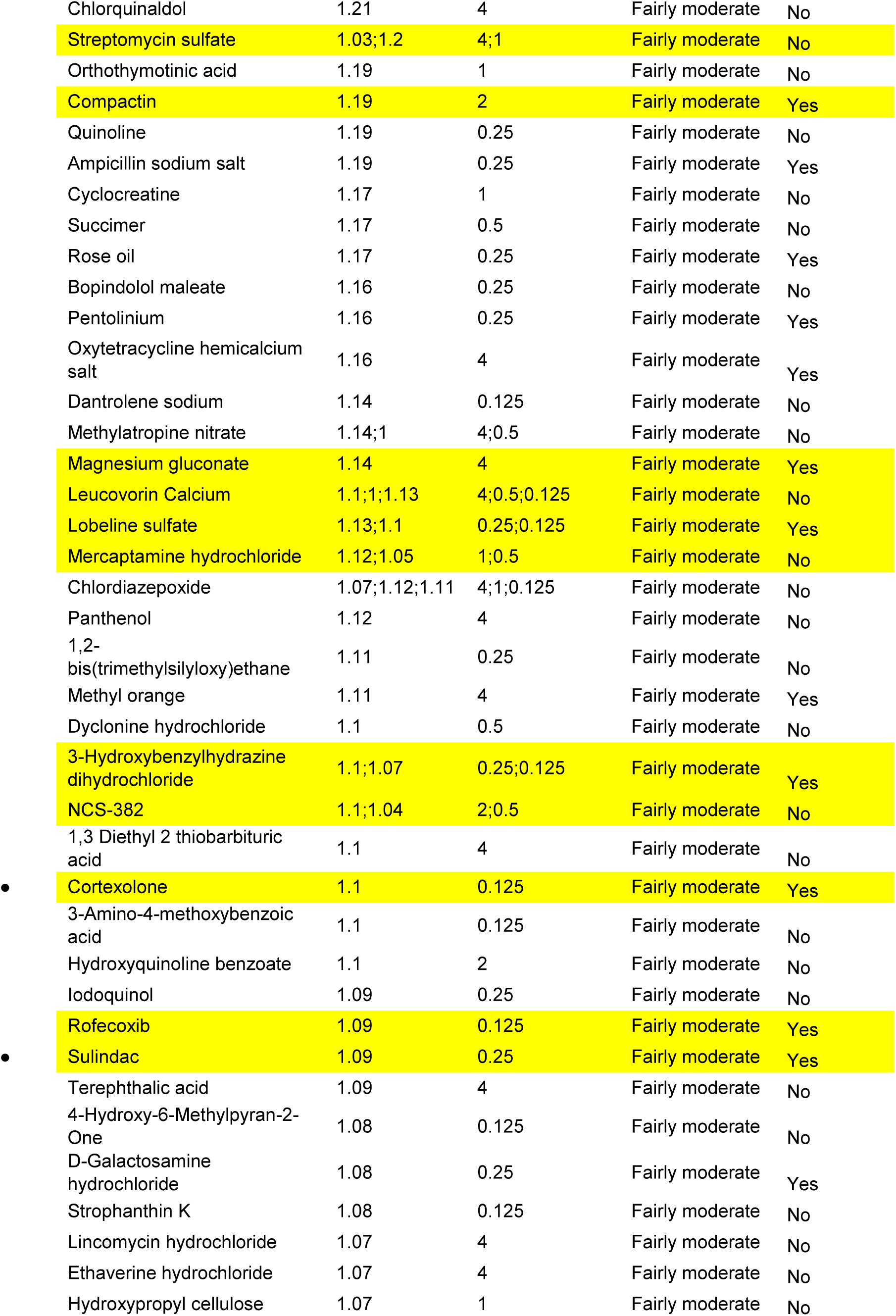
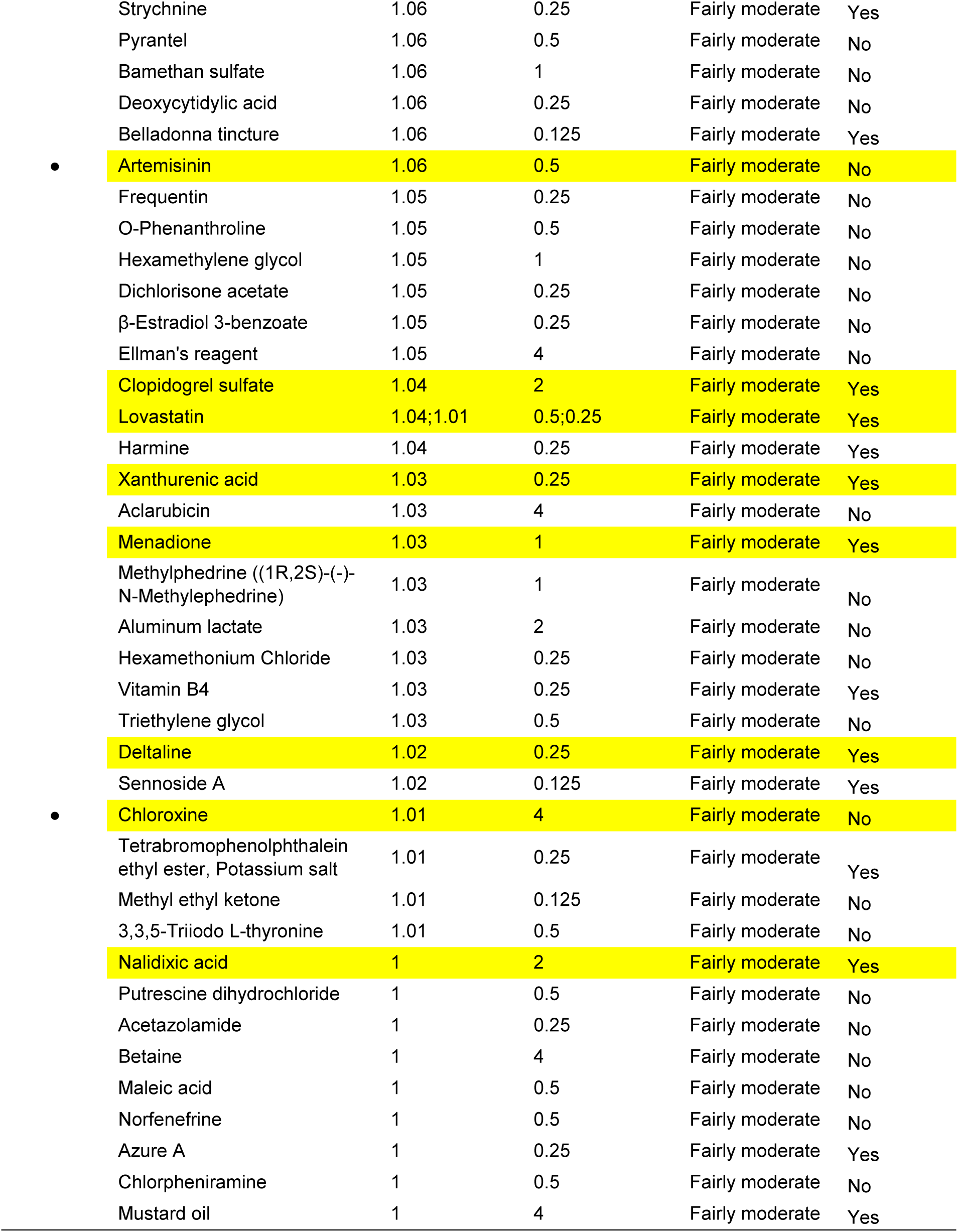
The 113 hit compounds from the primary screen ordered according to SSMD score. Compounds producing a SSMD of ≥1 were considered hits. Drug names, concentrations producing SSMD ≥1, SSMD scores, effect subtypes, and dose-dependent trend results are shown. Yellow highlighted drugs were selected for the confirmation tests. “●” denotes validated lead compounds.

**Supplementary Table 4.**
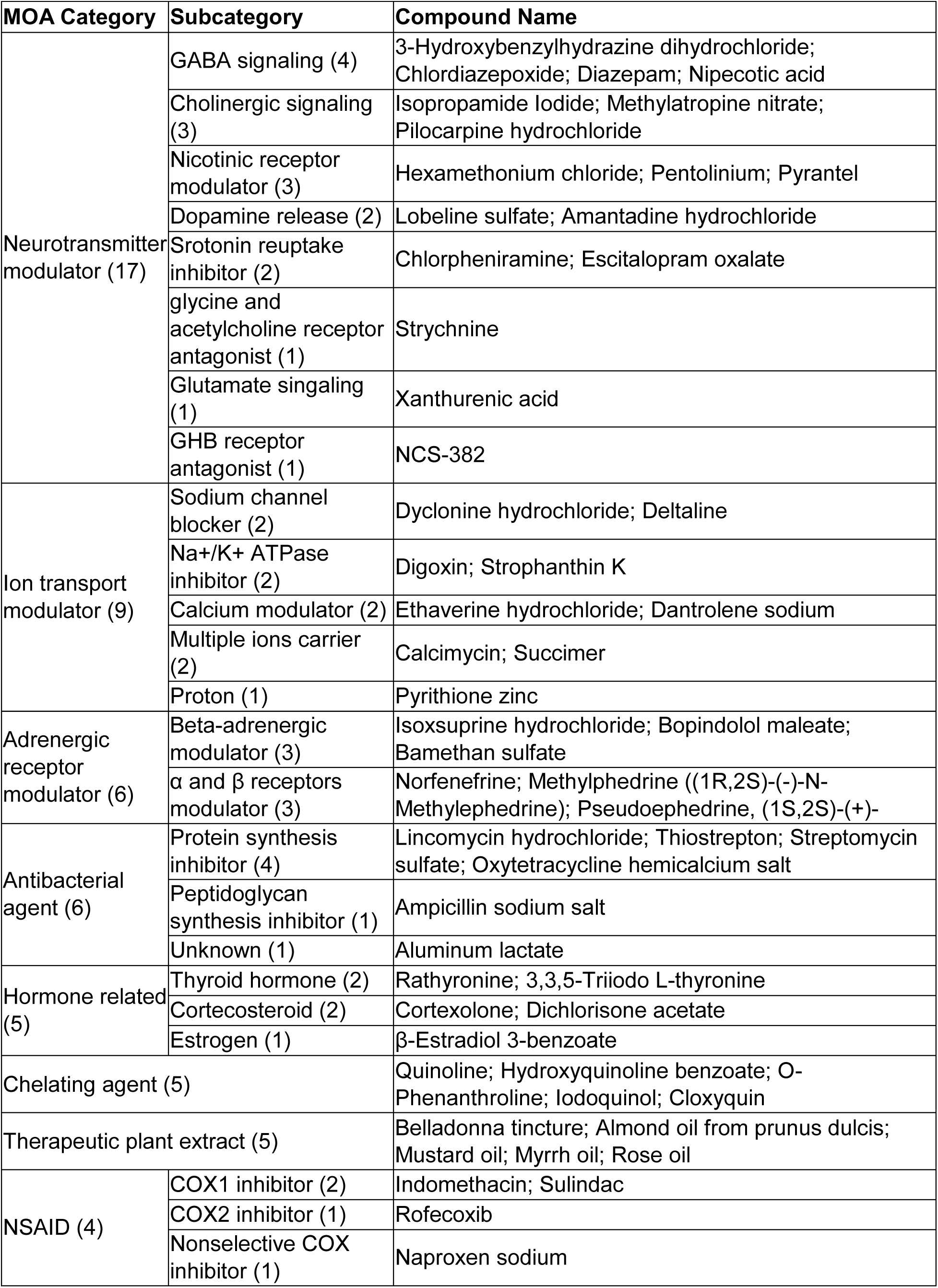
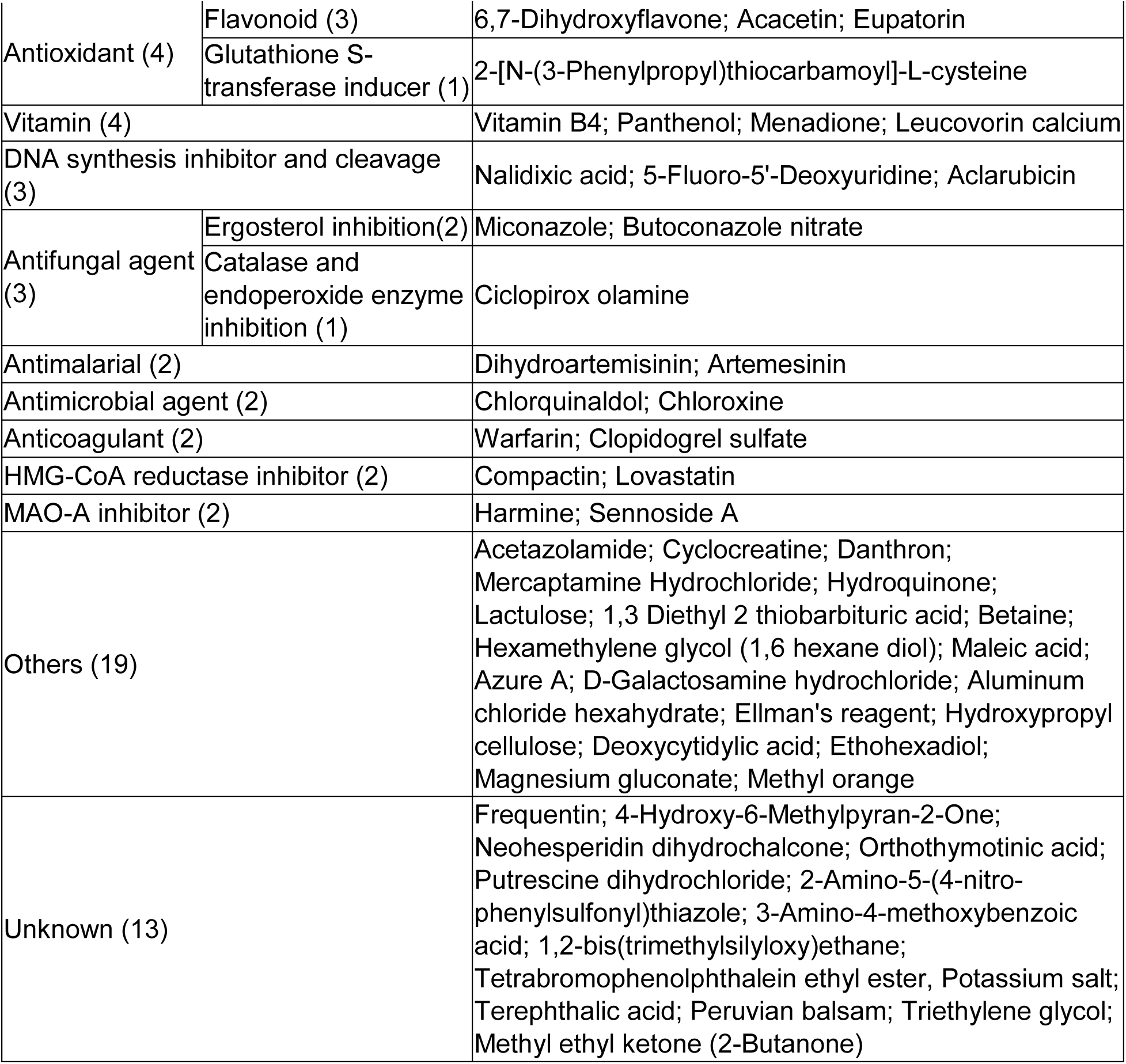
Clinical “on label” MOA of the 113 hit compounds from the primary screen. MOA are listed in order from most common to least common. The names and number of compounds in each category and subcategory are listed.

**Supplementary Table 5.**
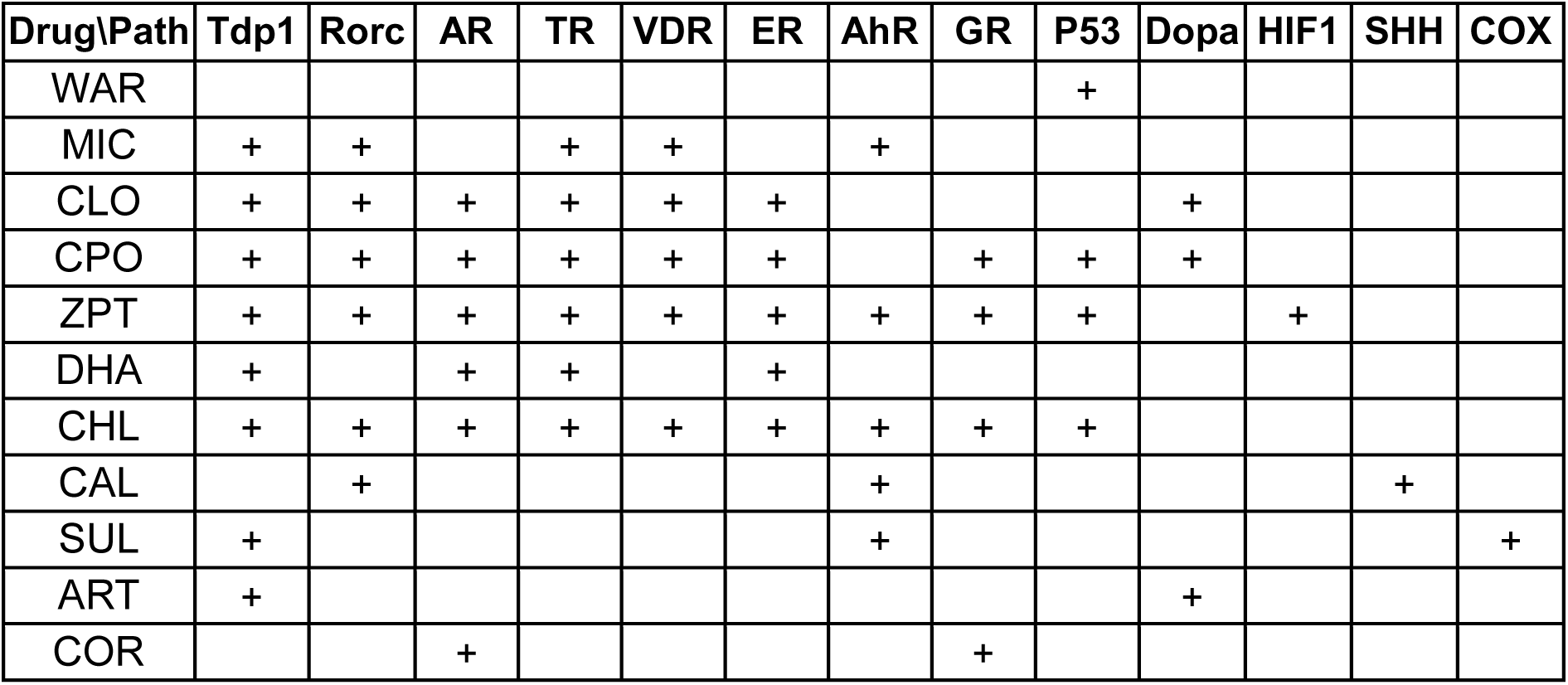
Summary of PubChem bioassay analysis. Results of thirteen target-based screens, with “+” indicating inhibitory activity. Targets: TDP1: Tyrosyl-DNA phosphodiesterase 1; Rorc: RAR-related orphan receptor gamma; AR: androgen receptor signaling pathway; TR: thyroid receptor signaling pathway; VDR: vitamin D receptor; ER: estrogen receptor alpha signaling; AhR: aryl hydrocarbon receptor signaling; GR: glucocorticoid receptor; Dopa: dopamine related; HIF1: Hypoxia-inducible factor 1-alpha; SHH: Sonic hedgehog; COX: cyclooxygenase.

| Drug\Path | Tdp1 | Rorc | AR | TR | VDR | ER | AhR | GR | P53 | Dopa | HIF1 | SHH | COX |
| --- | --- | --- | --- | --- | --- | --- | --- | --- | --- | --- | --- | --- | --- |
| WAR |  |  |  |  |  |  |  |  | + |  |  |  |  |
| MIC | + | + |  | + | + |  | + |  |  |  |  |  |  |
| CLO | + | + | + | + | + | + |  |  |  | + |  |  |  |
| CPO | + | + | + | + | + | + |  | + | + | + |  |  |  |
| ZPT | + | + | + | + | + | + | + | + | + |  | + |  |  |
| DHA | + |  | + | + |  | + |  |  |  |  |  |  |  |
| CHL | + | + | + | + | + | + | + | + | + |  |  |  |  |
| CAL |  | + |  |  |  |  | + |  |  |  |  | + |  |
| SUL | + |  |  |  |  |  | + |  |  |  |  |  | + |
| ART | + |  |  |  |  |  |  |  |  | + |  |  |  |
| COR |  |  | + |  |  |  |  | + |  |  |  |  |  |

**Supplementary Table 6.**
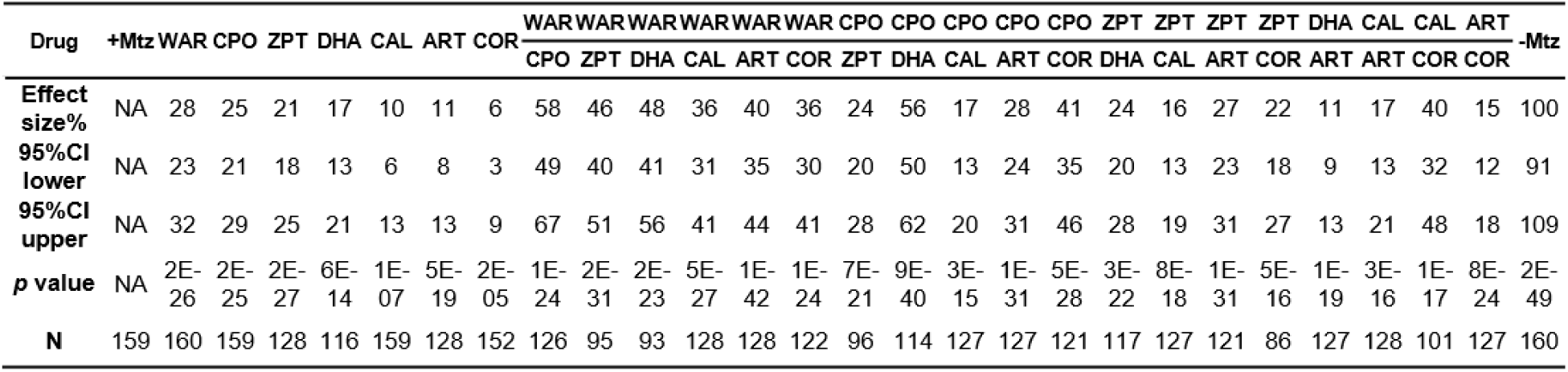
Statistical results of combinatorial assays (Figure 7). The effect size, 95% confidence intervals, *p* value, and sample size (N) for each condition is shown. *P* values were calculated by comparing each condition to +Mtz controls using Student’s *t* test followed by Bonferroni correction for multiple comparisons (α=0.0019 adjusted significance level).

